# Hybrid Quadrupole Mass Filter – Radial Ejection Linear Ion Trap and Intelligent Data Acquisition Enable Highly Multiplex Targeted Proteomics

**DOI:** 10.1101/2024.05.31.596848

**Authors:** Philip M. Remes, Cristina C. Jacob, Lilian R. Heil, Nicholas Shulman, Brendan X. MacLean, Michael J. MacCoss

## Abstract

Targeted mass spectrometry (MS) methods are powerful tools for selective and sensitive analysis of peptides identified by global discovery experiments. Selected reaction monitoring (SRM) is currently the most widely accepted MS method in the clinic, due to its reliability and analytical performance. However, due to limited throughput and the difficulty in setting up and analyzing large scale assays, SRM and parallel reaction monitoring (PRM) are typically used only for very refined assays of on the order of 100 targets or less. Here we introduce a new MS platform with a quadrupole mass filter, collision cell, linear ion trap architecture that has increased acquisition rates compared to the analogous hardware found in the Orbitrap™ Tribrid™ series instruments. The platform can target more analytes than existing SRM and PRM instruments – in the range of 5000 to 8000 peptides per hour. This capability for high multiplexing is enabled by acquisition rates of 70-100 Hz for peptide applications, and the incorporation of real-time chromatogram alignment that adjusts for retention time drift and enables narrow time scheduled acquisition windows. Finally, we describe a Skyline external software tool that implements the building of targeted methods based on data independent acquisition chromatogram libraries or unscheduled analysis of heavy labeled standards. We show that the platform delivers ~10x lower LOQs than traditional SRM analysis for a highly multiplex assay and also demonstrate how analytical figures of merit change while varying method duration with a constant number of analytes, or by keeping a constant time duration while varying the number of analytes.

## INTRODUCTION

Mass spectrometry-based proteomics has become an essential tool for biomedical research. Historically, most proteomics methods approach a new sample with little to no prior knowledge of the contents of the sample. Each experiment “discovers” peptides and proteins as if they have never been seen previously. The metric of success has frequently been the number of proteins or peptides that can be detected in a sample with little concern over the quality of the quantitative measurements. This often means that the instrumentation, methods, and approaches used may be less than optimal to detect quantitative differences because of multiple hypothesis testing and poor ion statistics.

The early ion trap instruments of the 80’s and 90’s were of the so-called 3D type, comprised of a ring and end cap electrodes and which had a spectral space charge limit^1^ on the order of 1000 ions. The 3D ion traps were popular for the first wave of proteomics applications where the experimental emphasis was largely qualitative. Quantitation, if performed, was often done with spectral counting^2^. The 3D ion traps were difficult to use for true quantitative applications because of limited dynamic range, limited speed, and because isolation was hampered by space charge effects in the limited volume of the ion trap. The 2D linear ion traps (LIT) solved the bulk of the volume-related space charge issues, having spectral space charge limits 15x higher than the 3D devices.^1^ The LTQ version of the LIT had a 2-fold symmetric X-dimension electrode displacement, or “stretch”, to reduce the effects of the field distortions of the ejection slots. The asymmetry between X and Y introduced an axial barrier to ion injection to the LIT and reduced the effectiveness of ion isolation during injection. The LTQ Velos trap^3^ introduced a 4-fold symmetric X- and Y-dimensional stretch and improved isolation during injection, which was sorely needed due to the advent of the higher flux ion sources being introduced in the late 2000’s. A remaining important inefficiency of the Velos trap was the fact that ion accumulation and analysis could not be performed at the same time, which limited the accumulation duty cycle to around 30% with an acquisition rate of 10 Hz. The Orbitrap Fusion Tribrid series instruments introduced the combination of quadrupole mass filter, collision cell, Orbitrap, and LIT, where finally the precursor isolation step could be performed essentially instantaneously, without any space charge issues, and pipelined ion accumulation and analysis processes allowed precursor ion utilization rates in the 80-90% range to be achieved^3^. For most Tribrid experiments the LIT has been used in its traditional role to generate spectra that can be used qualitatively for database or library searching^3^ as well as for simultaneous precursor selection in the data dependent tandem mass tag (TMT) sample multiplexing schemes^4^.

In the early 2000s in an effort to improve protein quantitation, there was a push to perform targeted experiments on specific proteins of interest within the sample^5^. These targeted mass spectrometry assays were viewed as a promising alternative to immunoassays in the clinical lab.^6,7^ There was significant development of tools and workflows resulting in targeted proteomics being acknowledged by Nature Methods in 2012 as Method of the Year.^8^ Those early methods were performed using an acquisition strategy known as selected reaction monitoring (SRM) on a triple quadrupole mass spectrometer. This was powerful as this hardware was common in mass spectrometry labs focused on development of targeted assays, and even today SRM-based assays are the gold standards for rigorous quantitative measurements, particularly in highly regulated environments. Since 2012, parallel reaction monitoring (PRM) has become an increasingly competitive alternative to SRM, as the entire product ion spectrum is measured, simplifying the method development workflow, enabling the user to confirm the identity of the analyte, to refine the product ion transitions that might contain interference post-acquisition, and actually providing a boost in sensitivity.

There has been nothing theoretically preventing the use of the Tribrid LIT for PRM experiments, except for the fact that PRM/SRM has traditionally been a precision tool for the analysis of a limited numbers of analytes. Furthermore, the resources of the Tribrid have been better deployed for upstream untargeted discovery studies. When the Tribrid has been used for PRM, the Orbitrap is used for mass analysis, largely because high resolution accurate mass analysis can simplify data analysis by reducing the incidence of interferences. However, recent work demonstrated that the LIT was more sensitive than the Orbitrap for targeted experiments, provided similar precision, and was not limited by selectivity if the transitions are refined after data collection^9^.

Despite the promise of targeted proteomics, there have been limitations in the number of peptides that can be measured due to the tradeoff between number of analytes and the sensitivity and precision of the measurement. In an effort to increase the number of targets that can be measured, data independent acquisition (DIA) has emerged as an alternative where relatively wide MS/MS isolation windows are acquired spanning a pre-defined mass range^10,11^. This acquisition intentionally samples many peptides in each MS/MS spectrum, decreasing the number of scan events necessary per cycle. This intentional multiplexing of the MS/MS acquisition using DIA comes at the expense of selectivity and dynamic range.

Another way to increase the number of peptide targets is to incorporate retention time scheduling into LC-MS/MS acquisition method. It is common to monitor each target for 2-5 min, despite peptide peak widths of <20 s. This inefficient scheduling is necessary because of chromatographic shifts that occur over the lifespan of an HPLC column. This irreproducible retention time is particularly pronounced in the nanoliter-microliter per minute flow rates. Recently we reported a prototype data acquisition strategy where the instrument automatically assessed the retention time relative to a reference analysis and adjusted the instrument target list to only sample a peptide in the time immediately prior and after the peptide was expected^12^. This was accomplished while accommodating non-linear and irregular shifts in retention time.

Another limitation of targeted proteomics has been that the development of targeted assays is complicated. This development often relies on the use of stochastically sampled protein digests using data dependent acquisition^13^ to determine which peptides should be targeted, followed by either the generation of synthetic peptide standards or production of recombinant proteins^14^ to optimize the method and parameters. Further logistical problems exist, such as the determination of which of the potentially tens of thousands of detected peptides are suitable for targeted analysis, whether they are still detectable on the targeted MS system, when they elute from the LC on a given day, and which precursor-to-product transitions should be used for SRM acquisition and analysis, or for PRM data analysis.

Recent progress was made in implementing and evaluating a streamlined strategy for building SRM assays from DIA gas-phase fractions (GPF), which are generated on a full-scan MS2 capable instrument^15^. In contrast to traditional wide-window DIA experiments^10,16^, GPF data are acquired with narrow isolation windows and longer injection times over several LC injections, essentially generating high quality PRM data on every detectable precursor. The resulting data, termed a chromatogram library^17^, is a rich source of information for redetecting analytes with targeted assays, containing retention times (RT), time-dependent extracted chromatograms and relative abundances for each potential fragment ion.

Here we introduce the Thermo Fisher Scientific™ Stellar™ mass spectrometer, which conceptually has the same hardware architecture as a Tribrid series instrument^3^ without the Orbitrap analyzer. In contrast to the earlier Thermo LIT-based instruments, the bulk of the development of the Stellar MS was focused on quantitative targeted MSn experiments, which are judged based on traditional analytical figures of merit like measurement precision, accuracy, limits of detection/quantification, ruggedness, and throughput. Along with hardware improvements that yield faster acquisition rates than previous generations of Thermo LITs, significant effort has also gone into the development of algorithms designed to improve targeted analysis through intelligent data acquisition. Chromatographic retention time shifts are accommodated with real-time updates to the scheduled target list with the methodology described previously^12^ and now termed Adaptive RT. Maximum injection times are dynamically adjusted based on the concurrency of the assays, yielding the largest possible injection times that respect a points-per-peak sampling rate requirement. Finally, software to aid in the rapid creation of targeted assays was developed to automate the GPF DIA to PRM strategy, which is embodied in a Skyline^18^ (ver.23.1.1.503) external tool called PRM Conductor. Skyline was used in this work to select LC peaks, integrate transitions and export results to external Python analysis scripts. The Stellar MS is a cost-effective platform for highly multiplexed targeted proteomics at an expanded scale relative to previous SRM and PRM technologies. Here we evaluate its performance in a set of three experiments designed to reveal the fundamental advantage of PRM over SRM analysis and the characteristic tradeoffs imposed by decisions about the LC gradient length and number of targets in the assay.

## EXPERIMENTAL PROCEDURES

### Sample Preparation

We prepared a matrix matched calibration curve using chicken plasma as a matched matrix for human plasma as described previously.^9^ Briefly, human and chicken plasma solutions were mixed in various ratios to keep a constant 333 ng/ul concentration. A 3x serial dilution was made in 7 levels from 50% to 0.2% human plasma with a 100% chicken blank for the final level. For the PQ500 experiments, PQ500 samples were obtained from Biognosys (Ki-3019-24). The samples were diluted at the manufacturer’s recommended level (120 μl diluent and then dilution by 4x). Digested human plasma was procured from Pierce (unreleased product in development) and used as a diluent at 300 ng/μl. A 3x serial dilution was created with 11 steps, from 1x manufacturer’s concentration=100% to 0.005% with a 100% plasma blank for the final level. For the *Esherichia coli* experiment, an *E. coli* digest was procured from Waters (PN 186003196) and HeLa digest from Pierce (PN 88329). The serial 2x dilution curve was 9 levels from 100 ng *E. coli* to 0.78 ng with a 200 ng/μl HeLa diluent, with a 100% HeLa blank for the final level. Pierce retention time calibration (PN 88320) (PRTC) mixture was added to all samples at 50 fmol/μl.

### Liquid chromatography

All experiments used a ES906A Easy Spray™ LC column heated to 45 C. The older human-chicken experiment used an RSLC-Ultimate 3000™ LC with direct injection, 1 μl/min flow rate, a 30-minute gradient and about 60-minute injection-to-injection time. The PQ500 experiment used a trap-and-elute injection scheme with 60 and 100 SPD methods on a Vanquish Neo™, that utilized 0.8 and 1.8 μl/min flow rates during the main portion of the gradient, respectively. The same Neo, column and 60 SPD method was used for the E. coli experiment. LC and instrument methods are described in more detail in the supplement.

### PRM Conductor Software

Skyline gives 3^rd^ party developers an interface to create programs referred to as “external tools” that can be launched from within Skyline to consume custom report data and extend Skyline’s capabilities^19^. PRM Conductor is an external tool that aids in the generation of targeted MS2 and MS3 assays. The creation of an assay is broken into 3 main parts. (1) Skyline metadata describing precursor-to-product transitions is used to filter transitions against a set of thresholds. The filters are absolute area, relative area, signal-to-background, correlation to median transition, LC base peak width, and retention time. Precursors are filtered by whether they have at least a certain number of qualifying transitions, typically 3. (2) Targeted assays are created with a visualization that depicts the concurrency of the assay in terms of how long the instrument would take to acquire data for the precursors as a function of retention time. Given a desired number of acquired points per LC peak and corresponding cycle time, the user can see how various experimental parameters affect the feasibility of acquiring data for the qualifying precursors, such as LIT scan rate and acquisition window width. (3) Finally, the user decides whether to create multiple assays for all qualifying precursors, or a single assay with a “balanced load” that picks the best N peptides per protein. Instrument methods for the final assay(s) can then be exported, based on a user-defined template method. PRM Conductor is included with the Stellar instrument control software and can also be freely downloaded from the Skyline Tools Store within Skyline. Tutorials can be found at the website, https://panoramaweb.org/prm_conductor.url.

### MS Instrumentation

The human-chicken experiment used a Thermo Fisher Scientific Altis Plus™ triple quadrupole mass spectrometer to compare with a prototype mass spectrometer, referred to as the Thermo Scientific Stellar Mass Spectrometer. Both instruments were calibrated prior to the experiment, and the same sample, column, and LC was moved from instrument to instrument to perform a fair comparison. The Altis has a traditional triple quadrupole design with two resolving quadrupoles, a collision cell and an electron multiplier, and is characterized by its very fast maximum acquisition rates of over 600 transitions/sec. The Altis and Stellar share much of the same basic instrument architecture, vacuum chamber and turbo pump (Figure 1). Both have a 1.6 mm x 0.58 mm letter-box style ion transfer tube. The Stellar has a newer, longer version of the ion funnel that was introduced with the Orbitrap Ascend™.^20^ This longer funnel accommodates a second, smaller ion transfer capillary to deliver calibrant ions to the system without user intervention, using a system called Auto-Ready™. The Stellar has the Q00 multipole with the low pass filtering capabilities introduced on the Orbitrap Lumos™ and Exploris™. The Stellar Q0 multipole has the same shape and neutral beam blocker as Altis, but adds the DC drag voltages of the other product lines. The Q1 mass filter is the 5.25 mm field radius device used on the Altis Plus, Orbitrap Eclipse, and Ascend™ instruments. The Stellar collision cell has the 90-degree bend of the Altis device, but with a version of the double split gate introduced on Orbitrap Fusion on the front that allows injection time linearity down to below 5 μs. The collision cell, like those of the Tribrid™ and Exploris™, serves the dual purpose of collisionally activating precursor molecules (called HCD in the instrument method editor) and storing the fragments while analysis happens in the LIT. This parallel processing architecture affords a high ion accumulation duty cycle of about 85% for a 65 Hz acquisition. The LIT uses the same fundamental structure introduced with the Velos Pro™ that has a 4.0 mm field radius and a 4-fold symmetric stretch of 0.76 mm^21^. The LIT has high- and low-pressure cells filled with Helium at estimated pressures of ~6 mTorr and 0.5 mTorr, respectively. The high-pressure region is optimized for receiving ions and can also be used to perform resonance collision induced dissociation (termed CID in the instrument method editor), and broadband waveform isolation, including the synchronous precursor selection (SPS) technique introduced on Orbitrap Fusion™. The low-pressure cell performs mass analysis at four standard scan rates of 33, 67, 125, and 200 kDa/sec, with typical full width half maximum peak widths of ~0.35, 0.5, 0.7, and 1.0 Th at m/z 622. The Stellar acquisition rates peak at about 175 Hz for small scan ranges, higher than the Ascend and Eclipse platforms which peak at about 100 and 75 Hz respectively. (Figure S1). More typical acquisition rates for peptide applications with scan range 200-1400 Th are 70 Hz, 43 Hz, and 43 Hz for the respective platforms. Smaller scan range widths of 600 Th or less can be practical if specific transitions of a precursor are known to be of high quality. For example, a particular peptide could have good transitions in the range 400-800 Th, and spectra could be acquired for that range at 125 Hz. This optimization is used for the 100 SPD PQ500 assay, which has the highest precursor concurrency, and the process of creating an instrument method with custom scan ranges is automated with the PRM Conductor program.

**Figure 1.**
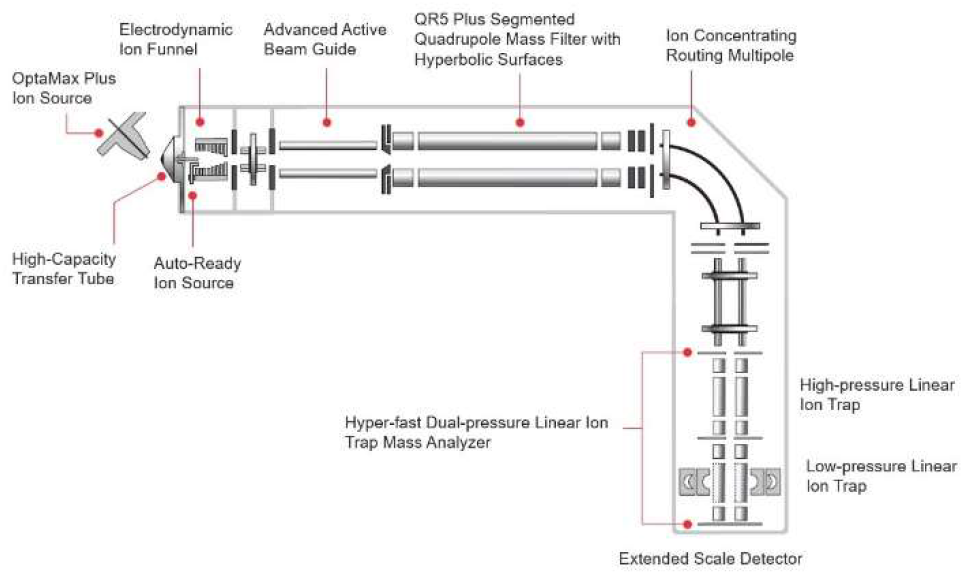
Instrument schematic of the Thermo Scientific Stellar mass spectrometer.

### Adaptive RT Algorithm

In SRM or PRM experiments, the period of time during which data are acquired for a target can be referred to as the acquisition window, which has a characteristic time duration, or width. We previously described an algorithm that adjusts for retention time shifts in real-time enabling scheduled acquisition windows of less than 1 minute.^12^ In brief, reference spectra from MS1 or DIA acquisitions can be compressed and serialized to a file having the extension .rtbin. This file can be selected and embedded into targeted MSn instrument methods, which causes the method editor to include those same MS or DIA acquisitions into the assay. During execution of the targeted MSn assay, the data from the MS or DIA acquisitions are compared with the embedded data in real-time to estimate retention time shifts compared to the reference and to update the set of active targets at a period of 3.5 times per LC peak width. Creation of the .rtbin file and embedding into a method can be automated with the PRM Conductor program, as described in the linked tutorials in the supplement.

### PQ500 PRM Assay Development

Neat PQ500 at the manufacturer’s recommended concentration was used to verify the retention times and presence of the standards. The Biognosys supplied transition list was imported into Skyline and an iRT calculatorsp was created from the supplied iRT values with PRTC selected as the reference compound list. Spectral library generation was performed with Prosit^22^ in Skyline. Unscheduled PRM methods were exported for both the proteom60 and 100 SPD methods that analyzed the 804 PQ500 compounds split into 10 fractions, where each subset also analyzed the PRTC compounds. The 10 fraction results were imported as a multi-injection replicate to Skyline, which used the iRT calculator and Prosit spectral library to aid in picking peaks. The PRM Conductor tool was used to create an instrument method for just the heavy peptides, to verify the retention times of the standards in 300 ng of plasma. Transitions were filtered without removing any PQ500 precursors. A final method was generated that included both the heavy and light peptides, and that included an Adaptive RT reference file for real-time chromatogram alignment. The 100 SPD method used 0.35-minute nominal acquisition window widths, while the 60 SPD method used nominal 0.6-minute widths. An optimization is applied in PRM Conductor that slightly widens the acquisition windows at the start and end of the runs. Additionally, the Adaptive RT algorithm applies a small buffer to the acquisition window of around an LC peak width, based on the uncertainty in the retention time shift estimations. See the supplementary materials for a walkthrough of the method generation process.

### Gas Phase Fractionation Library Generation

To generate targets for the human-chicken and E. coli-human studies, libraries were generated with the Stellar using 6 gas phase fractionation (GPF) DIA experiments having 1 Th isolation windows. The fractions each covered 100 Th precursor ranges, i.e. 400-500 Th, 500-600 Th, …, 900-1000 Th. The human-chicken library used 67 kDa/s analysis scan rate, while the *E. coli* library used 125 kDa/s. The data were searched in Proteome Discoverer with SEQUEST analysis for the human-chicken experiment, and with Chimerys analysis for the E. coli-human experiment. A normalized collision energy of 30% was used for all experiments, and a 200-1500 Th scan range. See the supplementary materials for raw files and instrument methods.

### Label-free PRM Assay Development

The Proteome Discoverer ver. 3.1.1.39 search results from the GPF libraries were imported into Skyline. The external tool PRM Conductor loads a report consisting of various metadata for each transition in the file. Transitions were filtered based on thresholds (vide supra). Precursors were accepted that had a minimum of 3 transitions meeting the requirements. As there were more accepted precursors than could be included in the PRM assay, multiple scheduled PRM assays were generated to further validate them by injecting them 3 times and filtering out those with a coefficient of variation (CV) > 30%. For the human-chicken experiment, a subset of 786 precursors was selected that had a maximum transition concurrency on the Altis of more than 600 transitions per second, using 2 min acquisition windows. For the E. coli experiment, two assays were generated, one with 397 precursors, and another with 1302 precursors. The E. coli assays used a nominal 0.6 min acquisition window. The human-chicken experiment was performed before any version of PRM Conductor was available, but the steps performed were the same as for the E. coli experiment, only using Python scripts.

### Points per Peak

Skyline computes each peptide’s acquisition points per peak as the number of acquisitions within its integration bounds. Skyline integration bounds are often wider than the actual width of the LC peak, and therefore points per peak are overestimated. To estimate points per peak we took the Skyline-determined full width at half maximum for each peptide, multiplied by 2.547 to convert to a base-width estimate for a Gaussian peak, and divided by the acquisition rate for each peptide. This more conservative estimate is about 1.5-1.6x smaller than the Skyline estimate. (Figure S2) See the supplementary information for a derivation of the 2.547 factor.

### Dilution Curve Analysis

The analysis of dilution curves is made difficult by the fact that, by design, the analytes of interest eventually are diluted below the limits of detection. Automated peak picking in Skyline currently is not informed by the results from more concentrated replicates, such that at lower concentrations the integration boundaries typically don’t match the high concentrations. A Python script was created to align the lower concentration replicates to the higher concentration replicates, outputting an integration boundary list that was imported back into Skyline. Limits of quantification and detection (LOQ and LOD) were determined offline with a similar script as was used previously^9^ which picks the set of transitions ≥ 3 transitions that give the lowest LOQ, even for the Altis data. The new script imposes that the LOQ be greater than 2x the LOD. This methodology was also described^23^ at ASMS but was not yet incorporated into an official Skyline release. Additional analyses were performed with Python scripts in Jupyter notebooks (see supplementary materials). For the PQ500 assay, the manufacturer supplied a spreadsheet of individual peptide amounts which allowed to transform the dilution levels from percent to peptide amount on column.

## RESULTS AND DISCUSSION

### SRM vs PRM for Plasma Analysis

The term parallel reaction monitoring (PRM) conveys the idea that all product ions of a precursor are formed and accumulated at the same time and mass analyzed in a single acquisition event. PRM has a fundamental advantage over the selected reaction monitoring (SRM) experiment, where each product ion is accumulated and analyzed in a set of serial events. All other things being equal, PRM has at least an N-fold advantage in sensitivity over SRM in the analysis of N products of a precursor, given that they all can be formed at the same collision energy. The PRM/SRM advantage grows even larger as injection or dwell times decrease and become significant compared to the overhead of SRM hardware switching for each product ion. The PRM/SRM sensitivity advantage is maximized for high throughput targeted proteomics, because precursors typically form many products, they can usually be formed at the same collision energy, multiple product analysis is typically desired to ensure sufficient selectivity^24^, and dwell times approach the 1 msec approximate hardware overhead times.

In the first experiment of this paper, we demonstrate the effect of this convergence of factors on analytical figures of merit in a comparison of SRM analysis on an Altis Plus to a prototype the Stellar mass spectrometer. An assay was created to target 786 plasma peptides in a 30-minute gradient with 2-minute acquisition windows. The Altis assay included up to 5 transitions per peptide. This assay pushes a triple quadrupole to its limits with a maximum of 624 concurrent transitions and a corresponding minimum dwell time per transition of 1.4 msec. (Figure S3) A PRM instrument like the Stellar, in contrast, has a maximum of 128 concurrent precursors for this assay, which receive 13 msec injection time per transition. With Adaptive RT real-time alignment, acquisition windows of ≤ 1 minute are feasible, which would have changed the maximum concurrent precursors to 68 with a minimum injection time 18.5 msec. The median ratio of injection/dwell times for PRM/SRM is 5.8 and 11.6 msec for the 2- and 1-minute acquisition windows, respectively (Figure S4). The effect of the increased accumulation time for the 2 min acquisition windows is evident in the figures of merit shown in Figure 2a and 2b, where 24% of peptides have LOQ < 20% dilution on the Altis, while 93% of peptides have LOQ < 20% for the Stellar. The LOD data are closer together, reflecting the fact that signal is decreasing with concentration appropriately for the Altis, but the number of ions is simply too small to achieve a high precision measurement under these conditions. These observations are illustrated in the specimen dilution curves of Figure 2c and 2d, and the histograms of relative figures of merit of Figure S5, where the median LOQ and LOD are respectively 15x and 2x better for the Stellar than the Altis.

**Figure 2.**
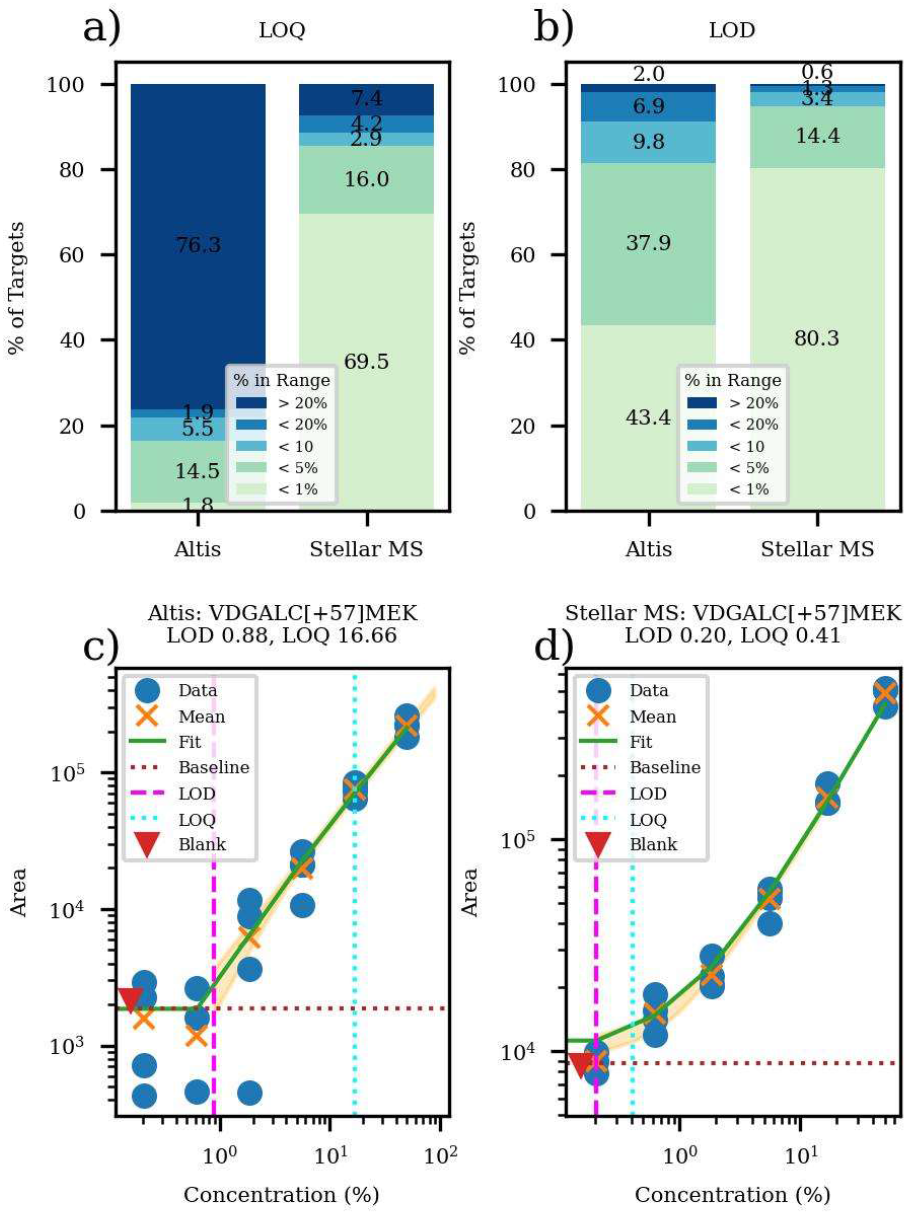
Figures of merit for matrix-matched human-chicken dilution with the Altis and the Stellar for 786 peptides. a) limits of quantitation, b) limits of detection and typical dilution curves for c) Altis and d) Stellar.

The Altis data in Figure 2 and the corresponding supplementary Figures S5 and S6 were optimized with the same script as the Stellar data, which chooses the ≥ 3 transitions that minimize the LOQ. The transition-optimizing procedure did not have much effect on the LOQ of the Altis data but did improve the LOD by 2x on average (Figure S6). In contrast, up to 15 transitions were considered for the full scan MS2 acquisitions and the LOQ optimization improved both LOQ and LOD by ~3x. In the Altis-SRM case, the unused transitions make the assay slower and less sensitive, which is not the case for the Stellar-PRM. This illustrates another fundamental advantage of PRM over SRM; transitions must be selected before running an SRM experiment, whereas post-acquisition transition selection effectively increases mass analysis selectivity for PRM and is particularly effective for peptide analysis where each compound usually has many viable transitions.

### PQ500 Analysis

The PQ500 kit is designed to perform absolute quantitation of ~578 plasma proteins. The kit consists of 804 peptides with C-terminal heavy labeling, and where the relative concentrations of the peptides are meant to reflect their abundance in plasma. This is a rather large set of peptides for this kind of experiment, considering that both the heavy and endogenous (light) versions of the peptides have to be analyzed to perform absolute quantitation. Creation of an assay is relatively straight-forward for a full scan MS2 instrument, however, because the LC peaks corresponding to the heavy labeled peptides can be easily found using the Prosit spectrum prediction capabilities built into Skyline and the iRT values supplied by Biognosys. With the Biognosys or Prosit calculated iRT’s, and with the high correspondence between predicted and experimental spectra (Figure S7), the identification of the correct LC peak was unambiguous in all but a few low intensity peptides. After identifying the heavy peptides and including both light and heavy peptides in one or more candidate assays, analysts typically perform a set of experiments to determine the assay feasibility. The analysis includes characterizing the number of acquired points per LC peak and the coefficients of variation of the measured peptide abundances. The 60 and 100 SPD assays had median points per peak of 7.1 and 6.8 respectively (Figure 3a), which is more than the generally accepted lower bound of 6 points which is approximately the minimum that meets the Nyquist criteria for a Gaussian peak. The coefficients of variation (CV’s) for 10 replicate injections these assays were 3.8 and 4.8%, respectively, with 94% of the peptides having CV less than the traditional 20% cutoff for both assays (Figure 3b).

**Figure 3.**
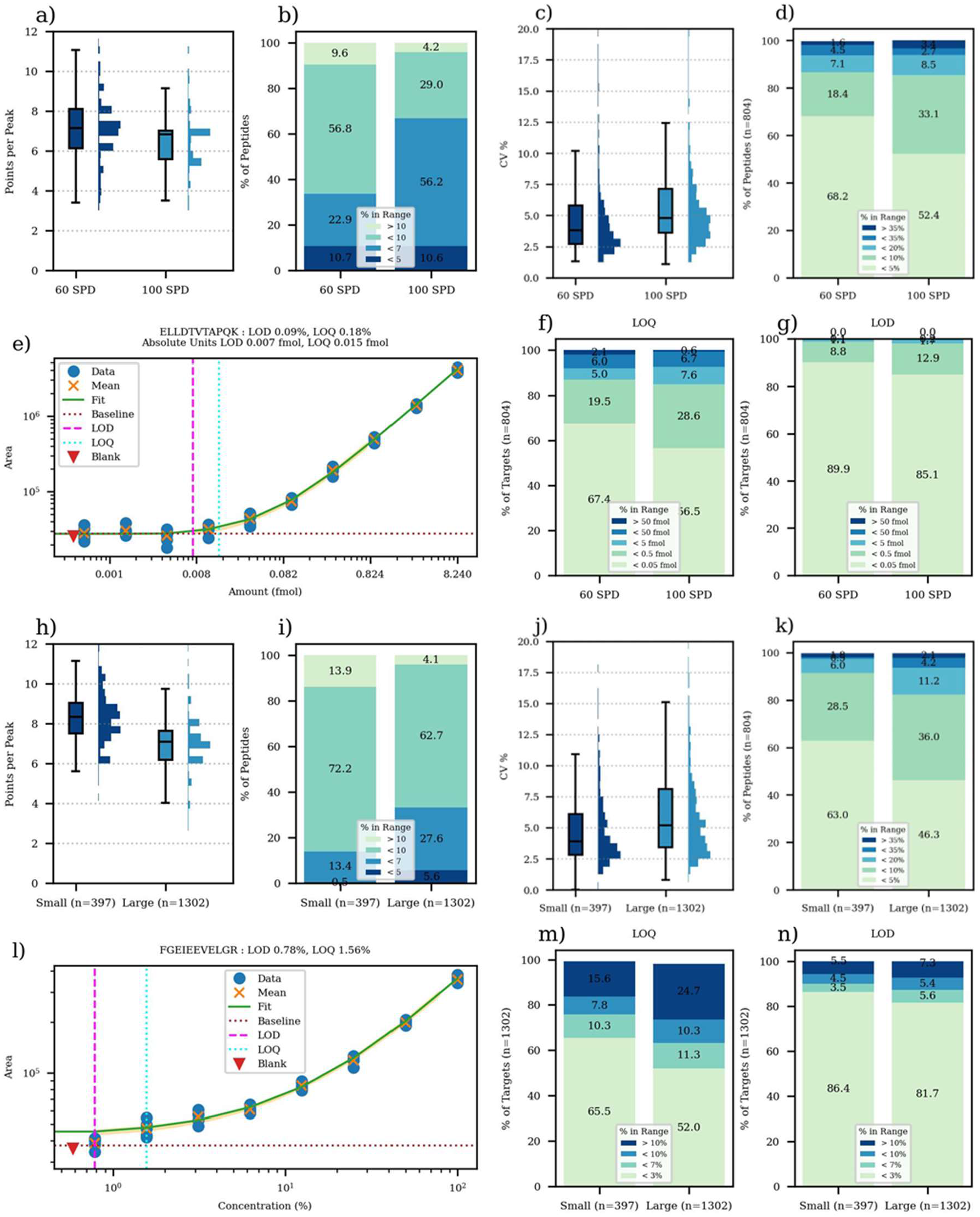
Summary of PQ500 study. a) Points per peak in box-whisker plot and b) stacked bar plots. c) CVs in box-whisker plot and d) stacked bar plot for 10 replicates. e) Specimen calibration curve for ELLDTVTAPQK peptide. f) Limits of quantitation and g) limits of detection. Summary of E. coli study. h) points per peak in box-whisker plot and i) stacked bar plots. j) CVs in box-whisker plot and k) stacked bar plot for 8 replicates. l) Specimen calibration curve for FGEIEEVELGR peptide. m) limits of quantitation and n) limits of detection.

**Figure 5.**
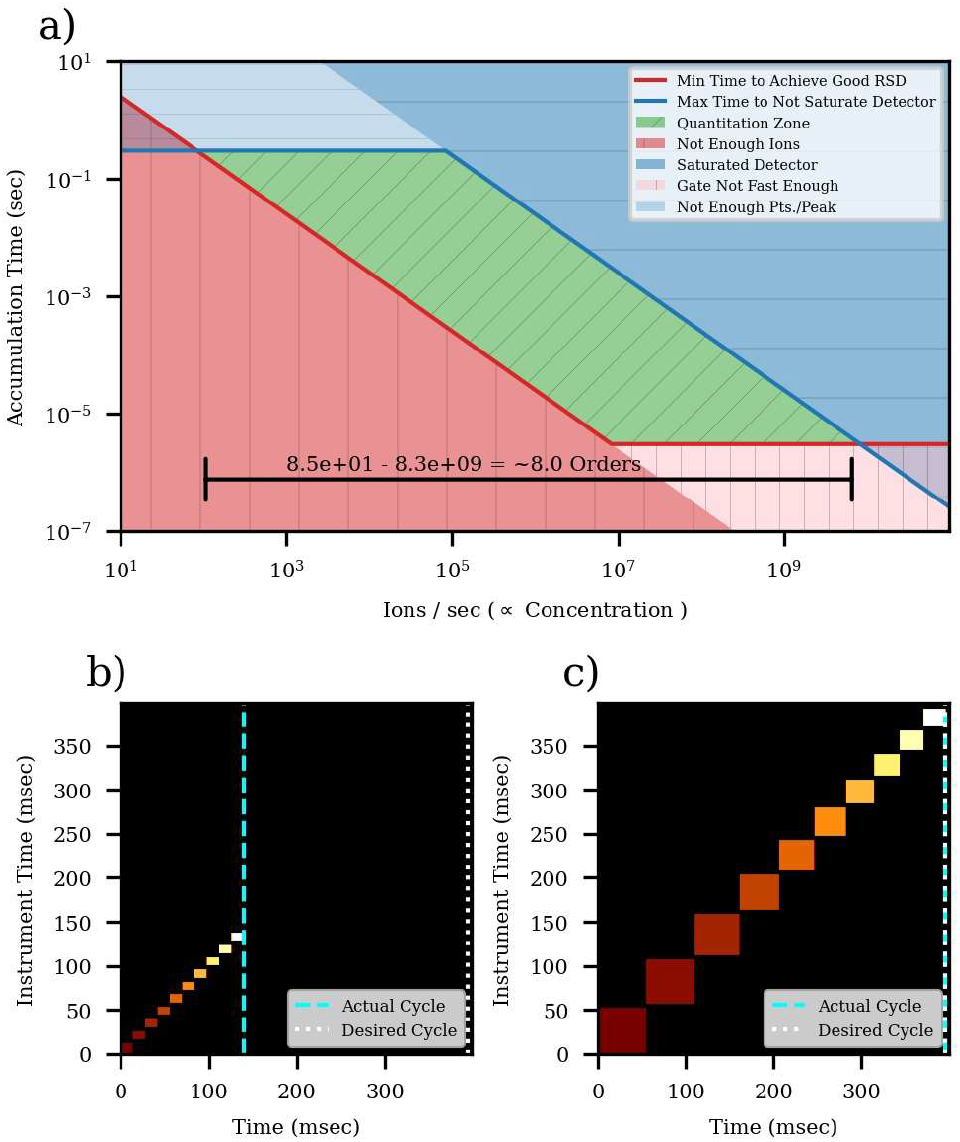
a) Theoretical depiction of how AGC expands the accessible quantitative range of concentrations. b) Assessment of active targets at the start of a cycle, with their minimum acquisition times on the y-axis, and where more intense precursors are white. c) Expansion of injection times inverse-proportionally to target abundance.

Dilution curves of the PQ500 into 300 ng of human plasma were performed to assess limits of detection and quantitation. A specimen calibration curve for a well-performing peptide ELLDTVTAPQK is shown in Figure 3c, which has an estimated LOQ of 15 attomoles. For both assays, the majority of peptides had an LOQ less than 50 attomoles, with about 85% of peptides having LOQ < 500 attomoles. The LOD’s were calculated as the bilinear turning points of the calibration curves and were nearly all less than 50 attomoles (Figure 3d). In terms of percent dilution, most LOQs for both assays were < 1% and 10% of LOQs were > 10% (Figure S8). Of interest to an experimenter is how the change in experiment time from 24 minutes to 14 minutes affects these analytical figures of merit, and Figure 3d gives an indication that the 60 SPD method is slightly better. Pair-wise comparison of the LOQs and LODs shows that both metrics are 1.6x lower for the 60 SPD method (Figure S9).

### E. coli Analysis

Label free targeted quantitation has 2x higher throughput than absolute quantitation because there are no heavy standards to measure for each compound. This type of analysis has traditionally not been favored as assays move closer to the clinic, where experiments are more conservative. The principal advantages of absolute quantitation are to normalize abundances for day-to-day instrument and LC drift, and to confirm the identity of the targets. While abundance normalization can be performed in less costly ways, such as normalization to the total ion current or to a small set of global standards, the problem of target confirmation has been more difficult. However, this conclusion is likely borne out of experience with SRM analysis, where streamlined assays may have only a single “quantifier” and “qualifier” transition for each peptide, and thus identity is confirmed with only the ratio of these transitions. With fast, full-scan MS2 analysis, peptide identity can be made more certain with traditional peptide sequencing tools, and label free quantitation could become more acceptable.

We created two label free 60 SPD assays for E. coli mixed with HeLa, by performing DIA GPF identification of possible targets and refining the results with PRM Conductor. The GPF results for 3 separate samples having 200 ng E. coli, a 1:1 mix of E. coli:HeLa, and 200 ng HeLa yielded 12.4k, 5.8k, 55.8k unique peptides, and 1.7k, 1.2k, and 7.1k protein groups respectively (Figure S10). Clearly the mixing of the two samples poses a challenge for peptide identification, so we started with the detections from the mixture sample to create two assays with 397 peptides (Small), and 1302 peptides (Large). Like for the PQ500 assay, replicate analysis for the Small and Large E. coli assays was performed, where the median points per peak were 8.3 and 7.1 and median CVs for 8 replicate injections were 3.9 and 5.2%, for Small and Large assays respectively (Figures 3h-k). For the E. coli assay, the lowest dilution level was 0.78%, which was too high to accurately assess the lower limits of many peptides, which have dilution curves like in Figure 3l. The majority of the peptides could be quantified below the 3% dilution level and nearly all below 10% dilution, sufficient for most real-world studies (Figure 3m-n). The Small assay LOQs were an average of 1.2x lower than the Large assay, while the LODs were nearly identical, although these results are likely influenced by the truncated dilution range at 0.78% preventing the reaching of the actual limits (Figure S11).

### Injection Time and Number of Ions

Thermo trapping instruments like the Stellar regulate the number of charges or ions per spectrum with a process called automatic gain control, or AGC. The basic concept is that a gate apparatus changes its transmission period (injection time) to pass an ideally constant number of ions per spectrum, and that dividing the detected abundance by the injection times yields a value proportional to ion flux in units of charges/second. As depicted in Figure 5a, the ability to vary the injection time expands the dynamic range of quantitation, from a base range given by just the storage and detector saturation limits and multiplies it by the gate linearity range. In the case of the Stellar, the pulsed nature of ion trap mass analsys^25^ limits the intra-scan dynamic range to about 3 orders (From 1 to 25k ions in a peak), while the gate can give a linear response from about 3e-6 seconds up to a practical experimental limit, for example 3e-1 seconds. Figure 5a was constructed by that the minimum number of ions required to achieve an RSD of 20%, due purely to counting statistics, is 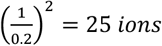^26^. Thus, the plot is separated into a red zone where ion flux is too low to achieve 25 ions, a blue zone where the detector would saturate with too many ions in a peak, and a green zone where precise quantitation could be performed. Practical dynamic ranges of quantification are subject to factors such as chemical background and ionization suppression, but not by ion storage capacity and detector saturation.

**Figure 5.**
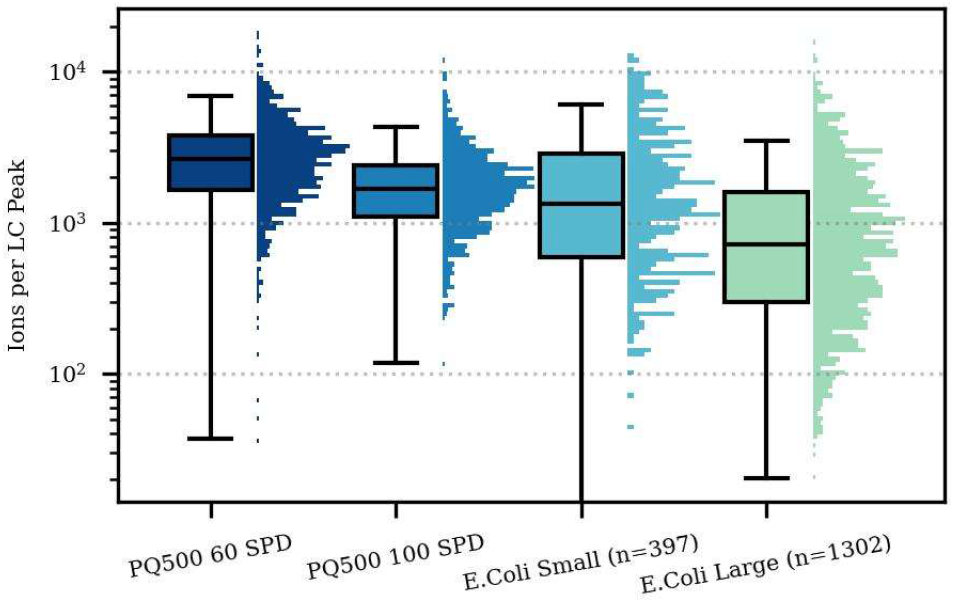
Number of integrated ions per LC peak for transitions belonging to each peptide, for the various assays. The target number of ions per spectrum was 1e4.

The AGC algorithms have evolved with time and reduce the number of parameters the user has to change. For targeted quantitation, an important development was the use of a “dynamic” maximum injection time. In real-time, at the beginning of a cycle of the active targets of a scheduled assay, the mass analysis-limited-time needed to acquire those targets is estimated and compared to the cycle time, as in Figure 5b. The remaining time in the cycle is distributed to the injection time of the targets inverse-proportionally to their abundance, so that less abundant targets receive more injection time, as in Figure 5c.

The injection times used for the 60 SPD assay are slightly higher than the 60 SPD assay and the Small E. coli assay injection times are higher than the Large assay (Figure S12). In both cases this is due to the lower concurrency of the 60 SPD and Small assays than their counterparts, which allows Dynamic AGC to use more of the extra cycle time for injecting ions. The concurrency of the assays is high enough to make SRM analysis quite difficult to impossible with a traditional 2 minute acquisition window, even if only using 3 transitions per precursor. The maximum number of concurrent transitions would be 759, 1281, 207, and 660 for the 60 SPD, 100 SPD, Small, and Large assays, respectively, while for PRM analysis the median concurrent precursors with the acquisition windows used in this study are 125, 139, 40, and 75 (Figure S13). The actual median acquisition window widths were 0.68, 0.4, 1.1, and 0.73 minutes, with nominal widths of 0.6, 0.35, 0.6 and 0.6 minutes (Figure S14). If the SRM experiment had extremely stable chromatography or a real-time alignment algorithm enabling the use these smaller acquisition windows, the median concurrent transitions would fall to 375, 417, 120 and 225. The relative comparison between SRM and PRM with Adaptive RT are still similar to the comparison for the Altis / Stellar experiment in this work, suggesting that the SRM instrument would be similarly challenged for the PQ500 and E. coli assays.

The number of ions per spectrum can be estimated by multiplying the spectral TIC by the injection time in seconds, assuming a charge state of 1+, which is reasonable for bottom-up MS/MS spectra. Care must be taken when using this procedure on different Thermo instruments, as Orbitrap intensities are scaled by an additional ~10x and for MS2 they may include other scaling factors that adjust for quadrupole transmission, for historical reasons. Estimating the actual number of ions can be useful because counting statistics give a lower bound on the ions required for a precise measurement. If the total integrated ions over an LC peak is less than 25, the precision of the measurement is unlikely to be better than 20%. In reality, other sources of variability, like the LC separation, electrospray ionization, the electron multiplication process, and chemical background raise the number of ions needed for a quality measurement. The number of ions from transitions of the peptide of interest in an LC peak at the LOQ and LOD was actually in the 200-500 range, on the order of 10x more than the theoretical lower bound (Figure S15). A simpler analysis of the number of ions in the replicate studies versus their CV shows the expected trend that high CV values occur predominantly for peptides with low measured numbers of ions (Figure S16). The trend is more pronounced for the E. coli peptides, which are present at their natural abundances than the PQ500 heavy standards which are spiked into the sample. The injection times at early and late retention times are sometimes larger because there are fewer active targets and Dynamic AGC allows to increase the injection time. The 60 SPD and Small assays receive slightly more injection time which explains the slightly higher number of ions (Figure 5), better CV and better quantitative metrics than their 100 SPD and Large assay counterparts (Figures 3).

## SUMMARY AND CONCLUSIONS

We tested the Stellar mass spectrometer in three different highly multiplexed targeted proteomics experiments. A ~10-fold improvement in LOQ and ~2-fold improvement in LOD was demonstrated compared to a triple quadrupole SRM instrument operated with dwell times that approach its transition switching times, demonstrating the fundamental sensitivity advantage of parallel compared to serial fragment accumulation. Reducing the experimental throughput from 24 to 14 minutes reduced the limits of quantitation by a factor of ~1.6x for a plasma assay, while increasing the number of targets from 397 to 1302 reduced the limits of quantitation by ~1.2x for an E. coli assay. These trends were explainable based on the changes in available injection time and number of ions that could be measured in the different scenarios.

Although these targeted experiments represent an advance in quantitative performance at high throughput compared to current SRM and PRM technology, they only utilize the MS2 HCD capabilities of the Stellar. The versatile LIT is also capable of resonance CID and multiplexed waveform-based isolation. These techniques enable SPS MS3 experiments, which offer a way to enhance selectivity at a cost of about 2x in acquisition rate. In future work, we plan to demonstrate how MS3 can be used judiciously to enhance the selectivity of a subset of analytes with unavoidable interference without a significant cost to the remaining analytes measured with MS2. Targeted SPS MS3 will also be used to analyze peptides labeled with tandem mass tags (TMT), increasing analytical throughput by up to potentially 10 times with the current TMTpro™ kits.

The software advances that enable the creation of these assays are meant to reduce or eliminate the use of spreadsheets and scripting. There is more that can be done to further automate and validate both method creation and data analysis. With more advances, we look forward to selective, sensitive targeted methods that rival DIA not only in terms of ease of setup and analysis, but also throughput and breadth of coverage.

## Supporting information

Supplementary LCMS Information

## ASSOCIATED CONTENT

### Supporting Information

Supplementary figures, Jupyter notebooks and metadata used for data analysis along with the RAW data and processed data (i.e. Skyline documents) from these experiments are available at Panorama Public (https://panoramaweb.org/stellar_ms_platform.url).

## ACKNOWLEDGMENT

This work was supported in part by National Institutes of Health grants R24GM141156, U19AG065156, and U01DK137097.

---

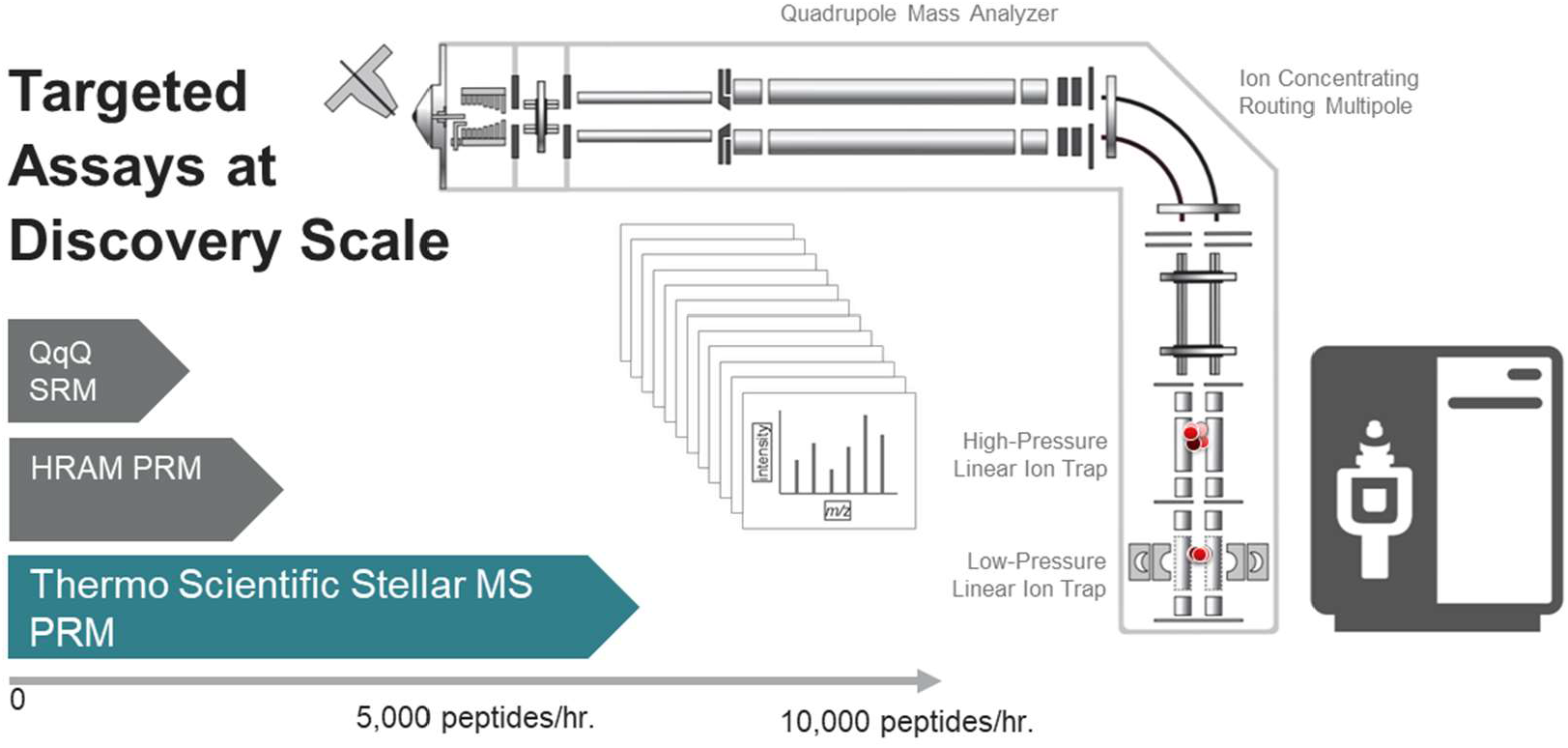

