## Supplementary LCMS Information for "Hybrid Quadrupole Mass Filter – Radial Ejection Linear Ion Trap and Intelligent Data Acquisition Enable Highly Multiplex Targeted Proteomics"

Supplementary LC/MS Information

A Vanquish Neo liquid chromatograph was used for the E. coli and PQ500 studies. The LC settings are given in the figure below.

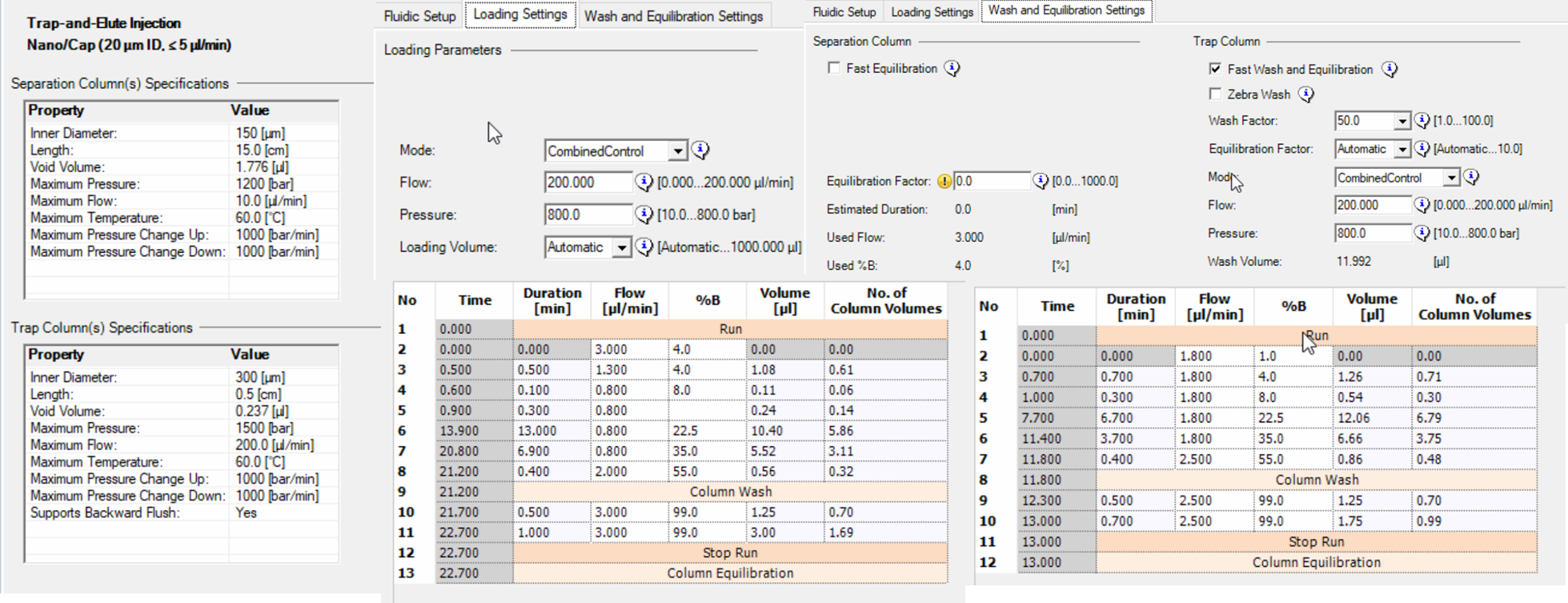

Screenshots of the targeted MS2 parameters for the 100 SPD and 60 SPD methods are given in the screenshot below. The scan range was 200-1500 for the 60 SPD methods, but was customized by PRM Conductor for the 100 SPD method.

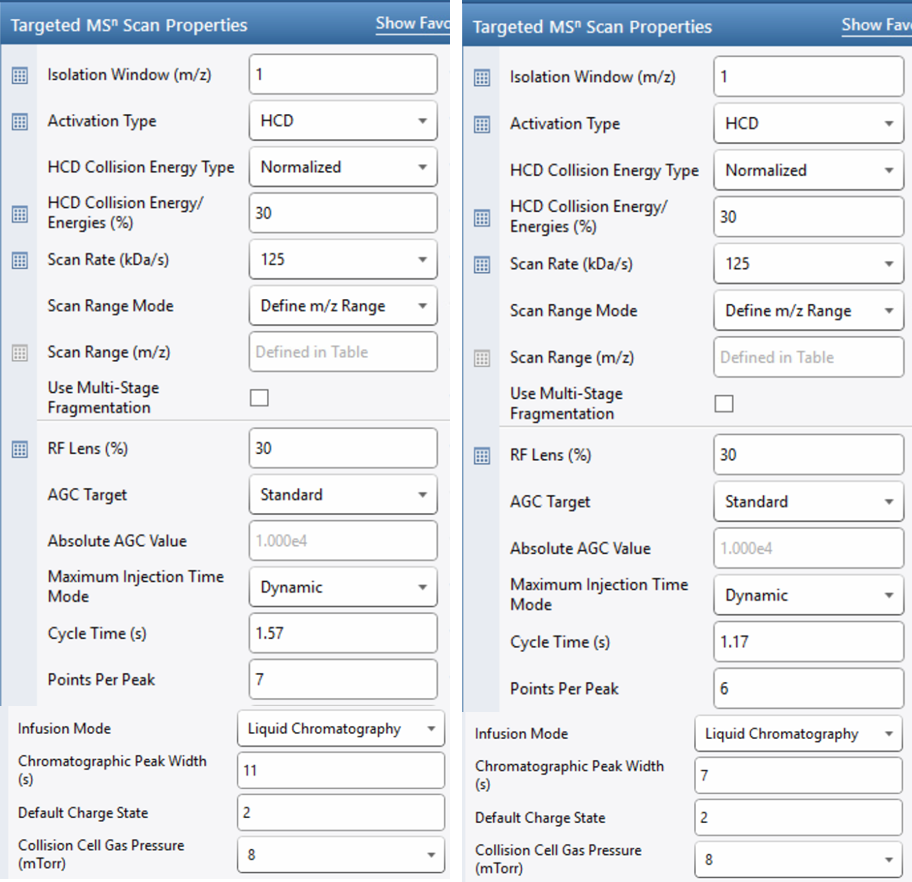
