## Supplementary Figures for "Hybrid Quadrupole Mass Filter – Radial Ejection Linear Ion Trap and Intelligent Data Acquisition Enable Highly Multiplex Targeted Proteomics"

**Conversion of Full Width Half Maximum Widths to Base Widths**

Consider a Gaussian peak where its full width half maximum (FWHM) is known. Substituting $x=\frac{FWHM}{2}$ in the equation of a Gaussian peak at half height $y=0.5$, and rearranging yields the following conversion between FWHM and the base width, defined here as 6 times the standard deviation.

$$0.5=\exp\left( -\frac{\left( \frac{FWHM}{2} \right)^{2}}{2\sigma^{2}} \right)$$

$$\ln0.5= -\frac{FWHM^{2}}{8\sigma^{2}}$$

$$\sigma=FWHM \left( \frac{-1}{8\ln0.5} \right)^{2}$$

$$BaseWidth=6\sigma=6 FWHM\left( \frac{-1}{8\ln0.5} \right)^{2}$$

$$BaseWidth=FWHM x 2.547$$

**Supplementary Figures**

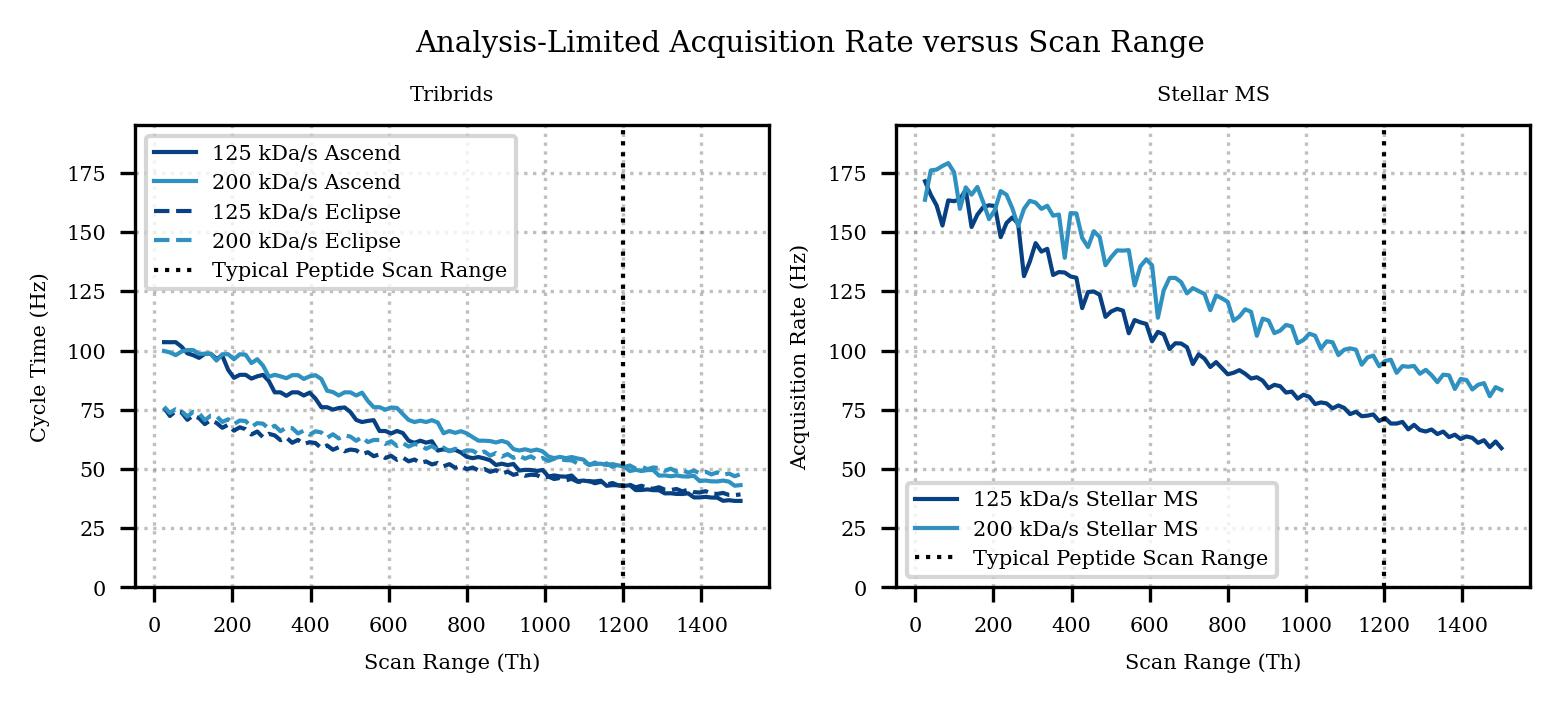

Figure S1. Analysis rate in Hertz versus acquisition scan range for the linear ion trap in a) Existing Tribrid platforms b) Stellar.

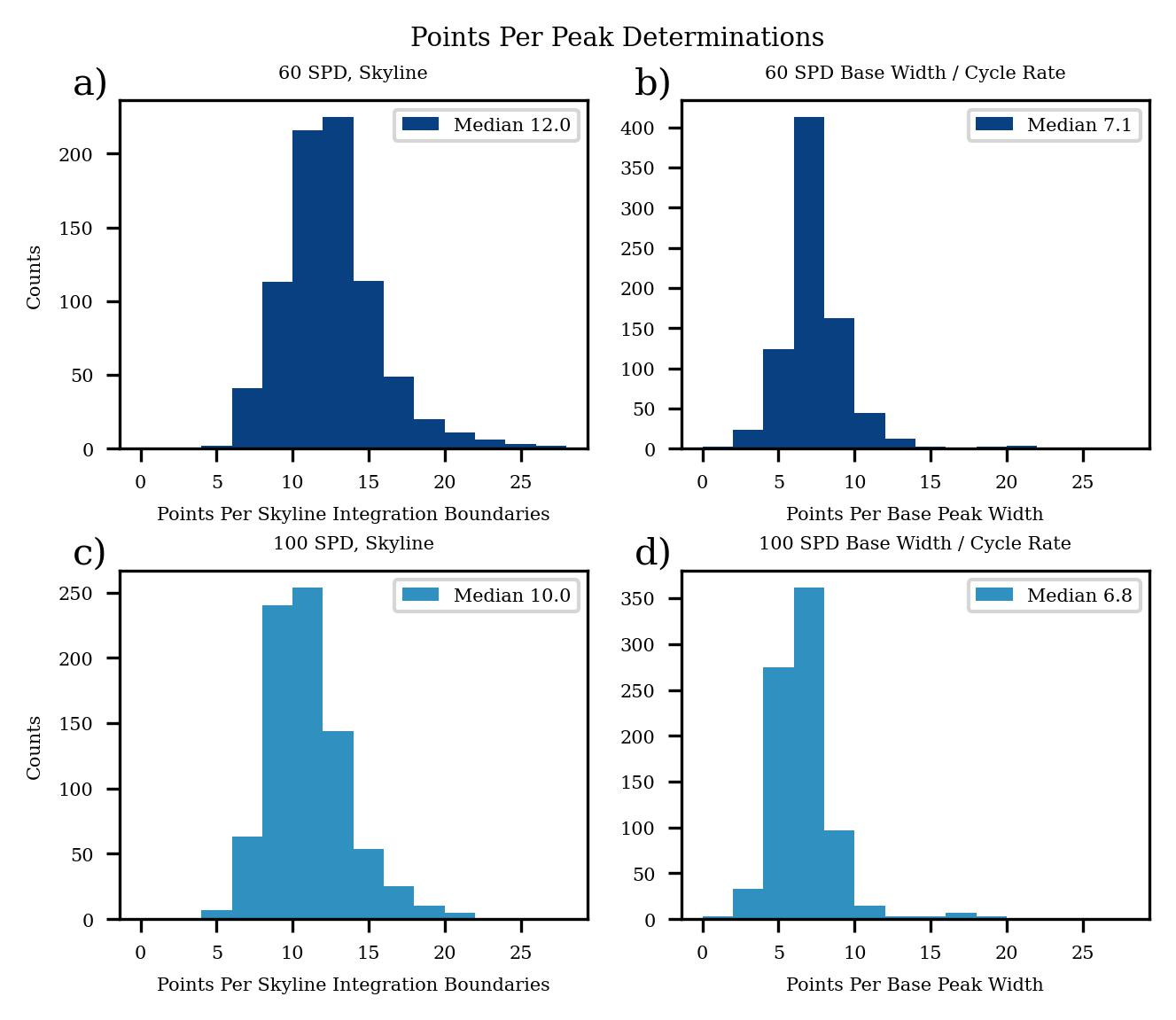

Figure S2. Histograms of PQ500 points per peak for 60 and 100 SPD using Skyline’s metric based on the integration bounds a,c) and based on an estimated base peak width in b,d).

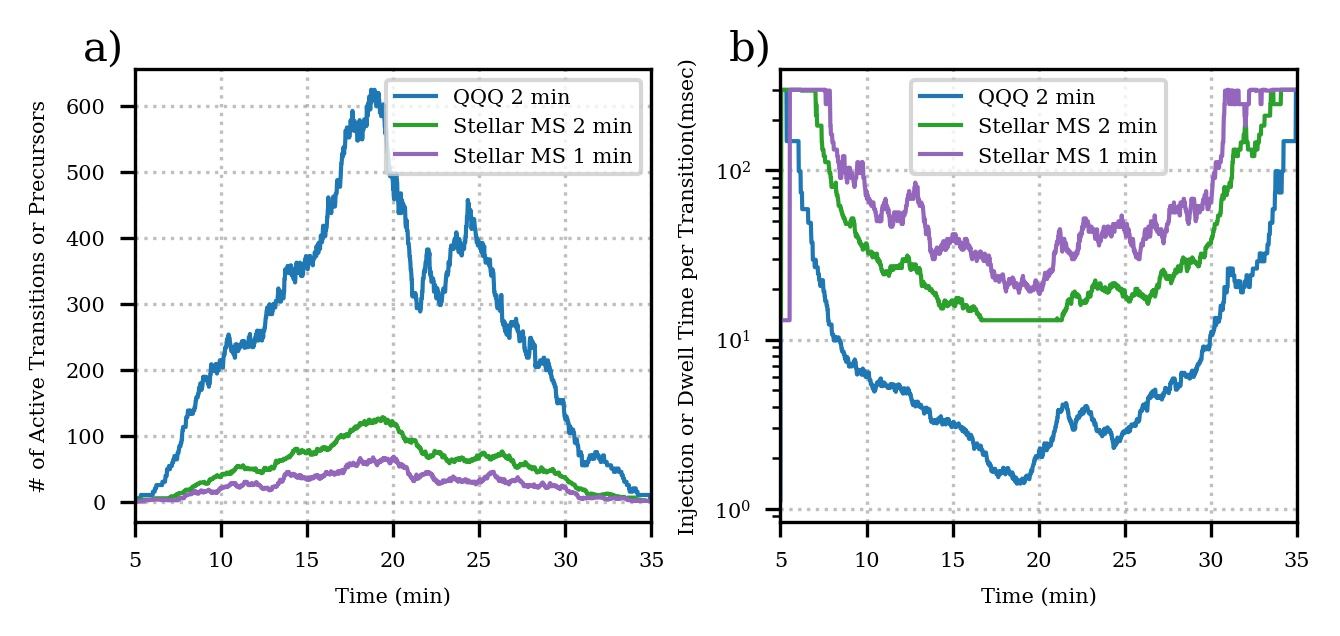

Figure S3. Metrics of the Human/Chicken assay for Altis and Stellar with 2 minute scheduled acquisition windows and an additional trace for a hypothetical 1 minute for Stellar. a) assay concurrency in active transitions for Altis and precursors for Stellar, b) dwell time for Altis and injection time for Stellar.

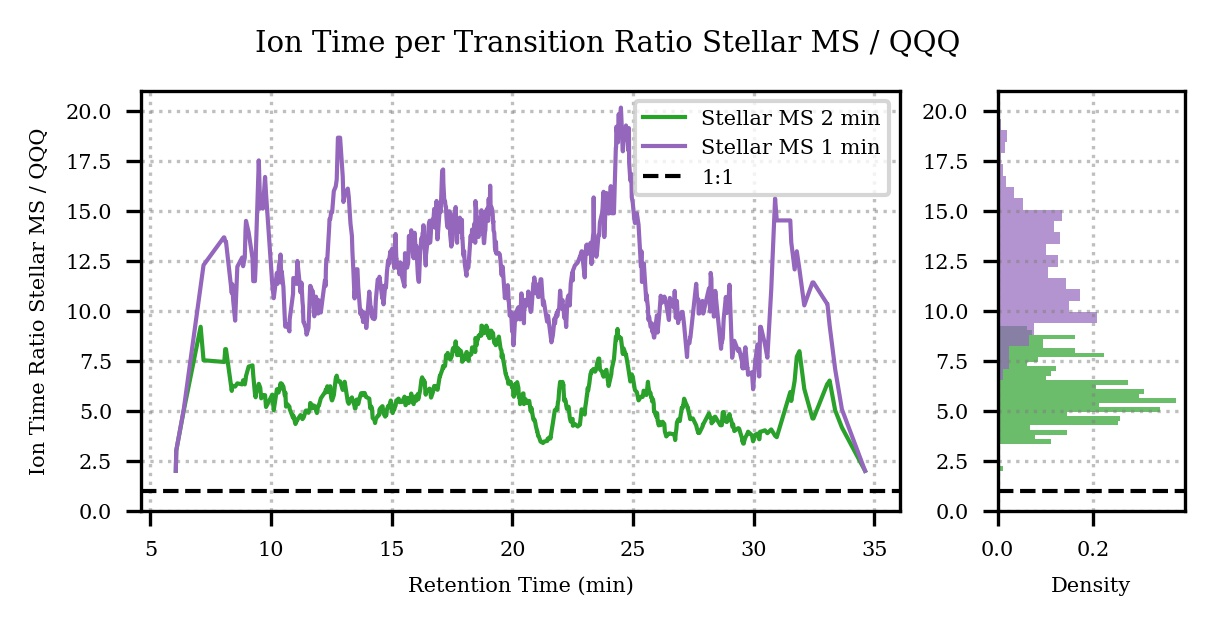

Figure S4. Ratio of injection time / dwell time per transition for Stellar / Altis, for the 2 minute and 1 minute data in Figure S3.

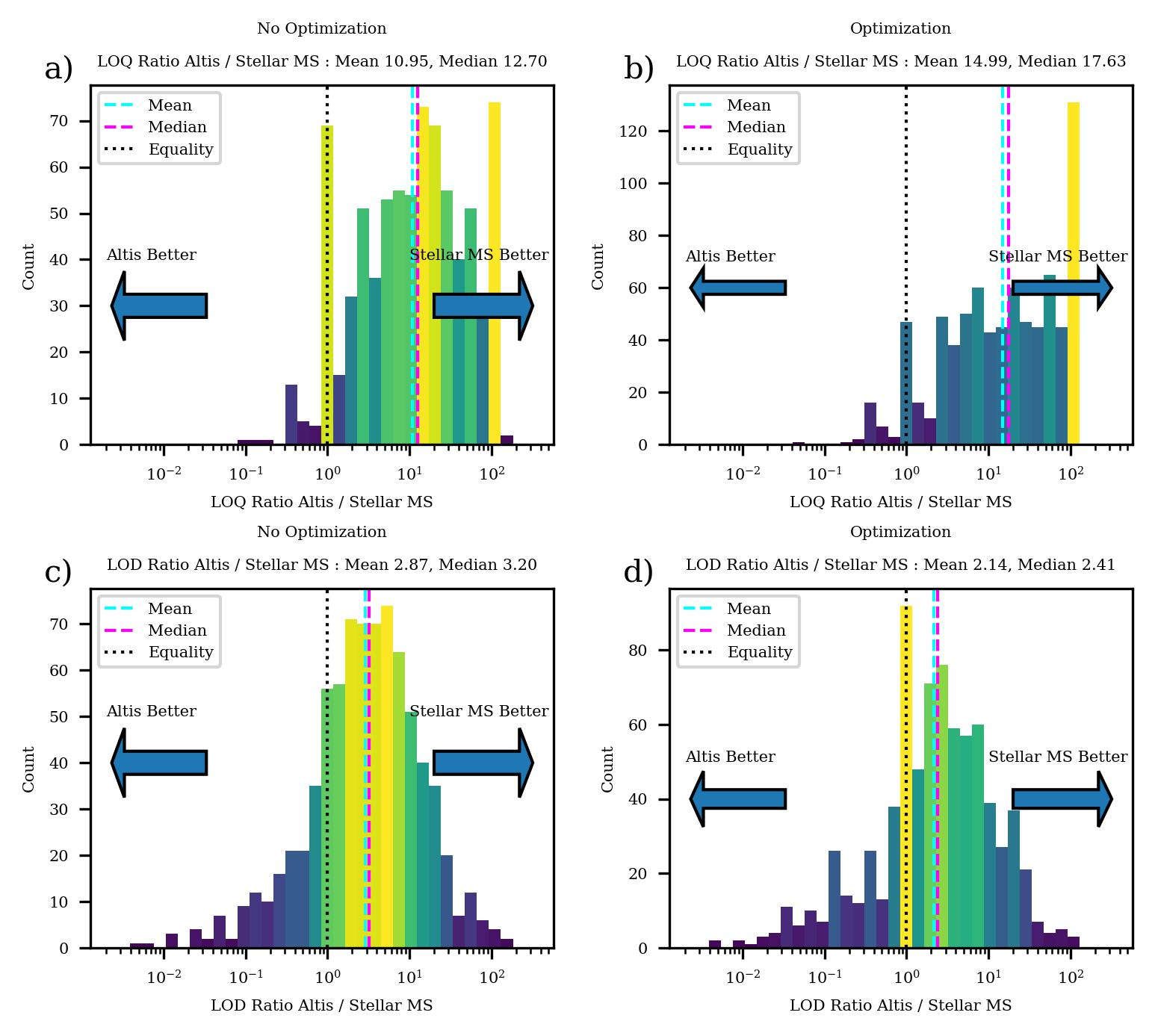

Figure S5. Comparison of figures of merit for the Human/Chicken dilution experiment. a) Ratios of Altis to Stellar LOQ’s without transitions optimization and b) with transition optimization. c) Ratios of Altis to Stellar LODs without transition optimization and d) with transition optimization.

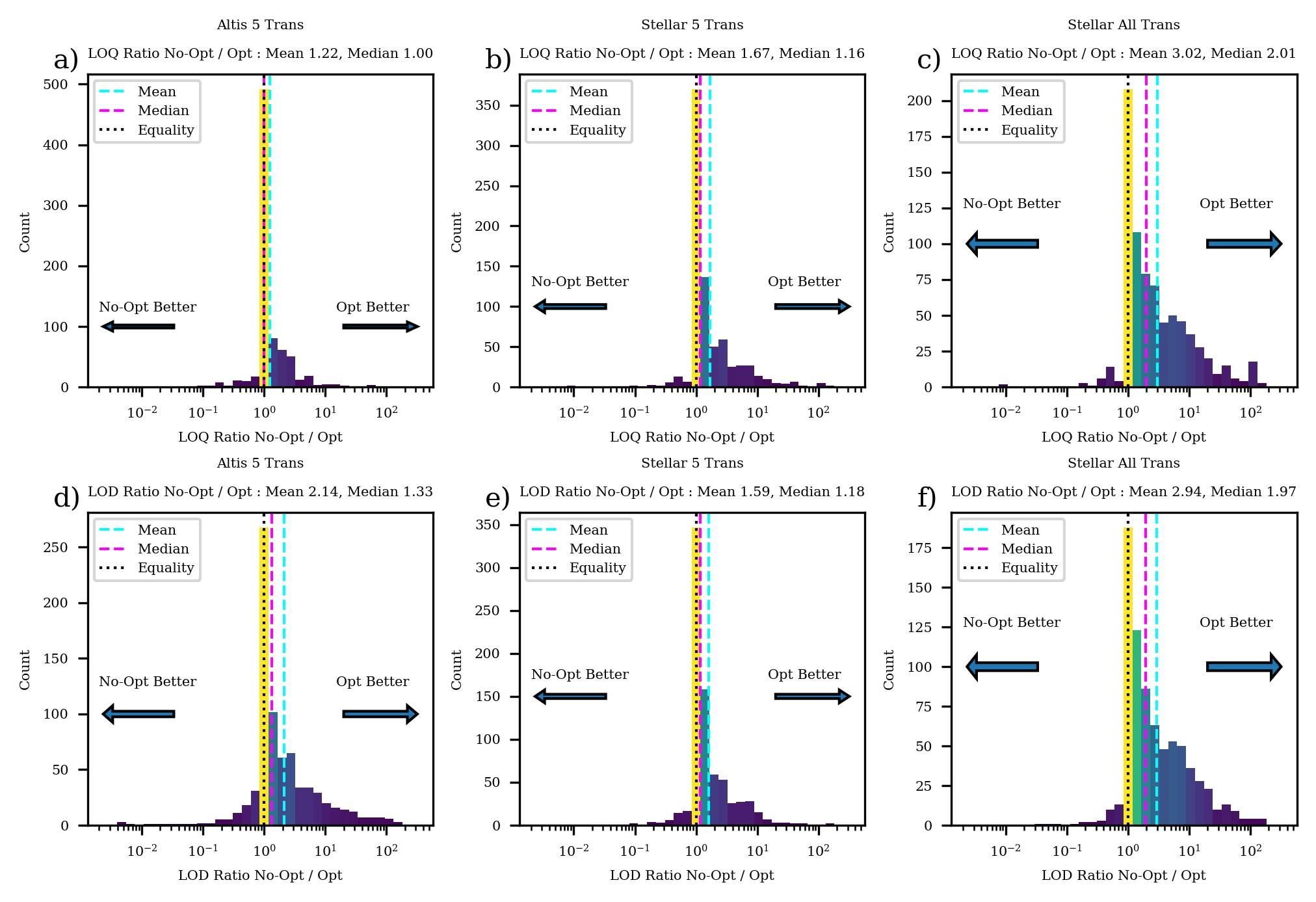

Figure S6. Comparison of figures of merit with and without transition optimization. LOQs are given for a) Altis top 5 transitions b) Stellar with the same top 5 transitions c) Stellar with up to 15 transitions. LODs are given for d) Altis top 5 transitions e) Stellar with the same top 5 transitions f) Stellar with up to 15 transitions.

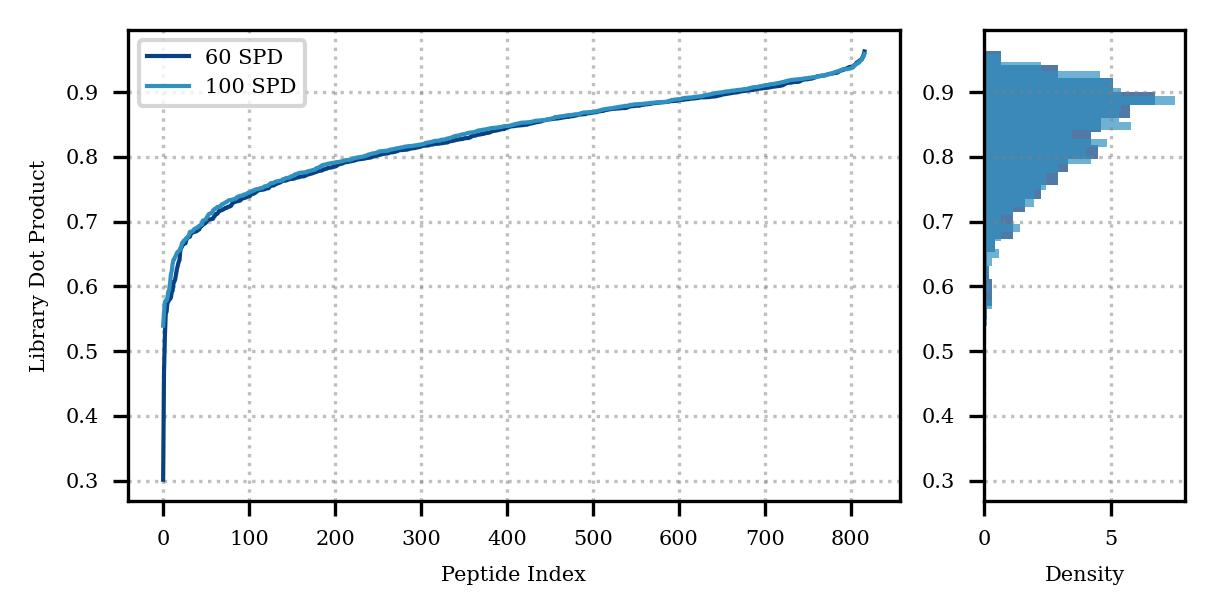

Figure S7. Distributions of the library dot product scores for the heavy PQ500 peptides compared to a spectral library generated with Prosit.

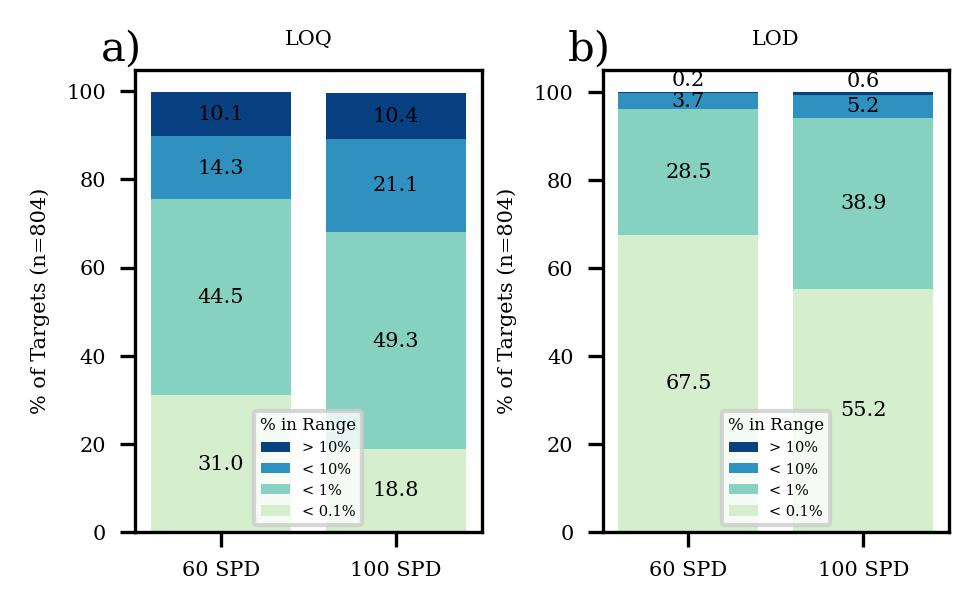

Figure S8. Distributions of figures of merit for the PQ500 assay in terms of percent dilution. a) LOQs, b) LODs.

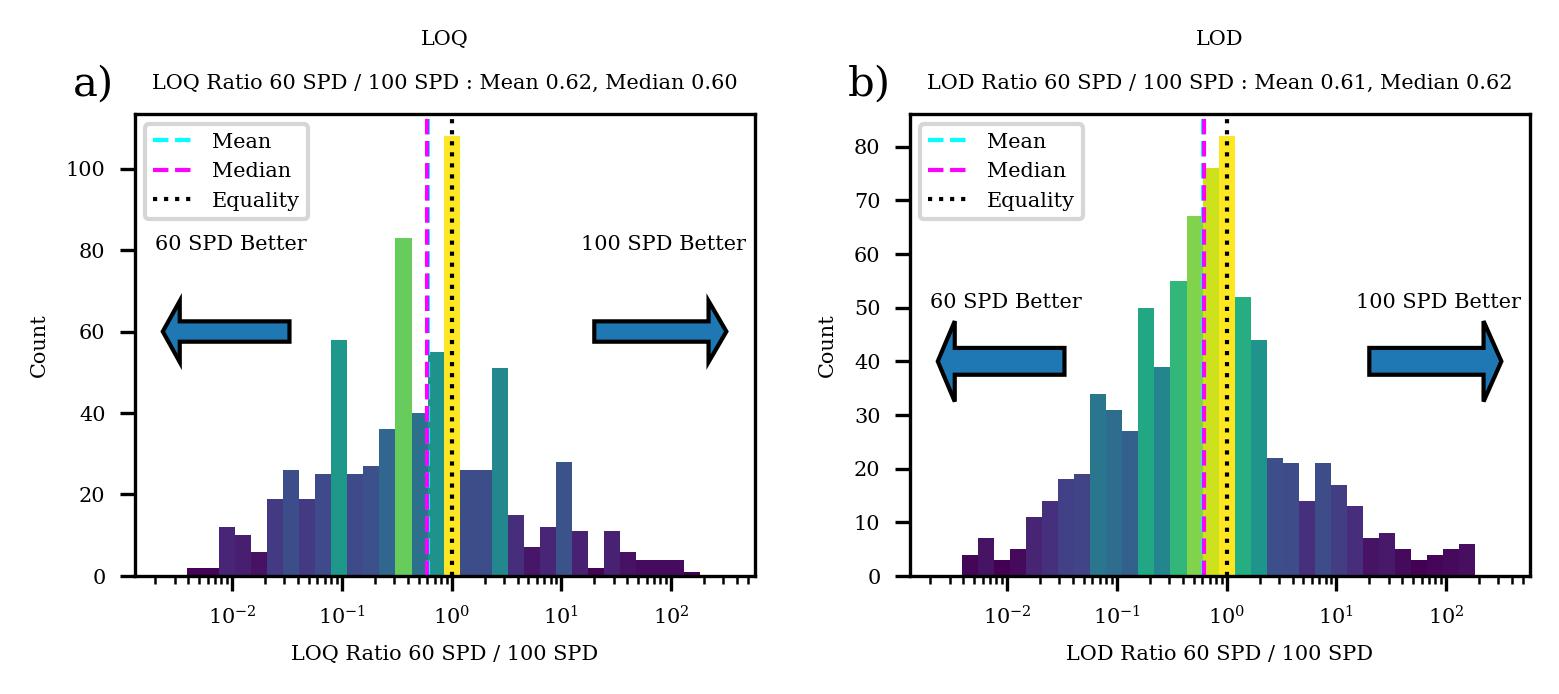

Figure S9. Comparison of figures of merit for the 60 and 100 SPD experiments of the PQ500 study. a) LOQs b) LODs.

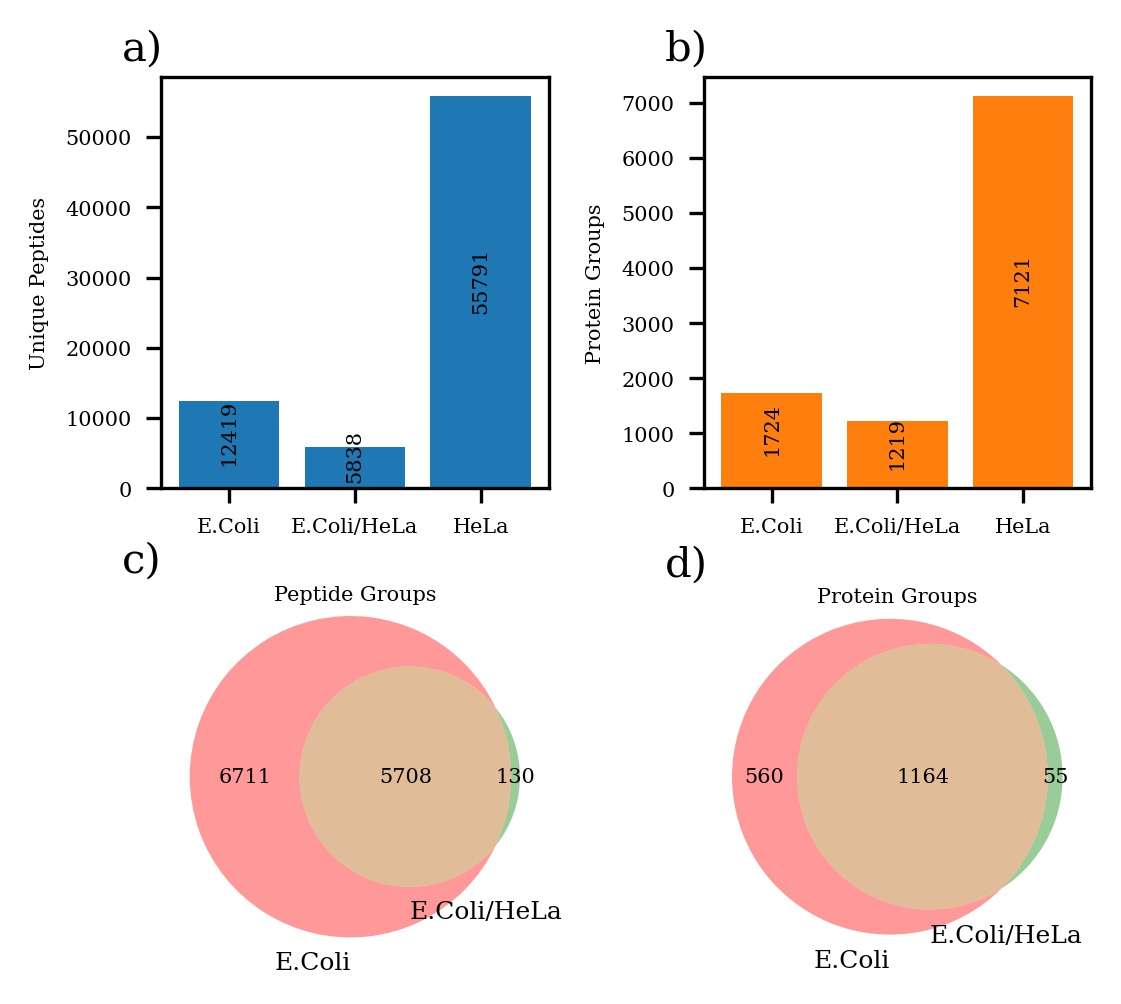

Figure S10. Summary of the DIA gas phase fractionation experiments of E. coli and HeLa.

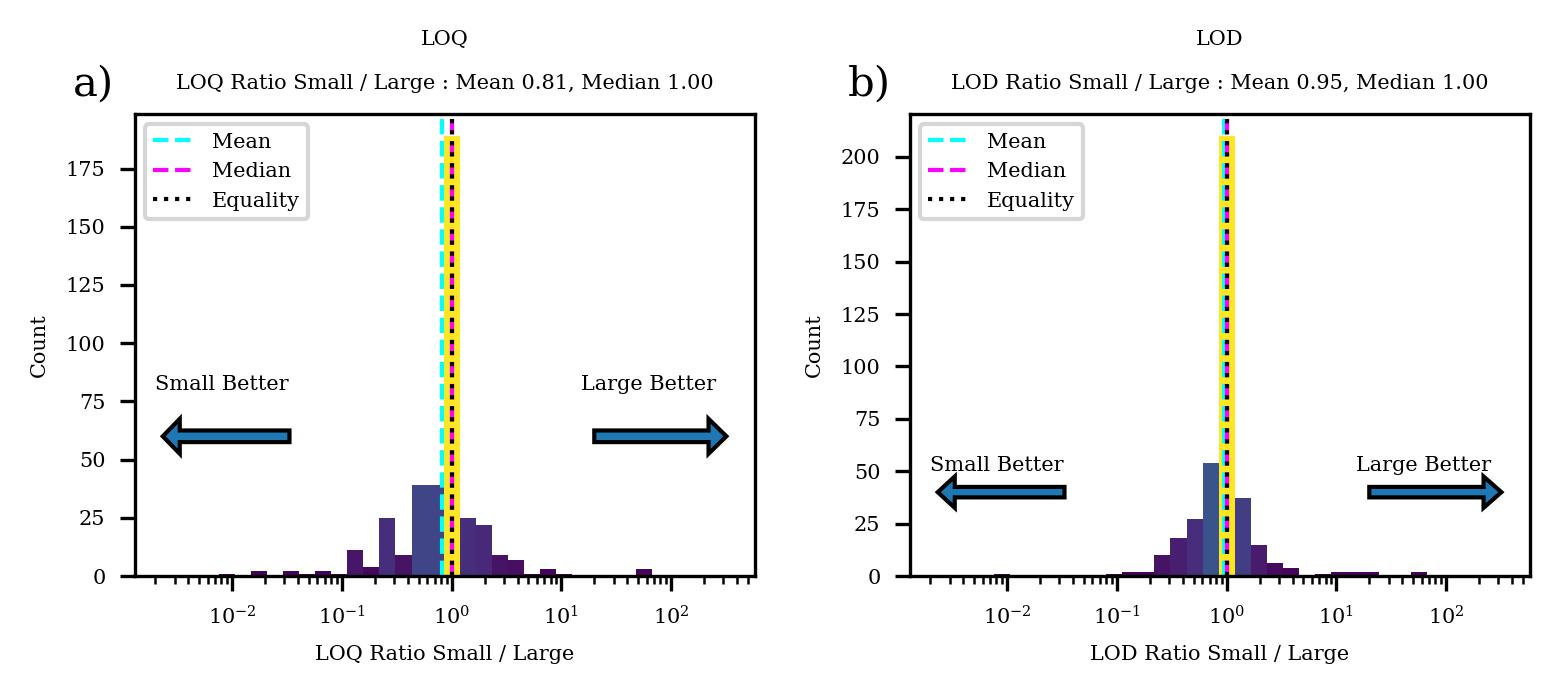

Figure S11. Comparison of figures of merit for the E. coli assay. a) LOQs, b) LODs.

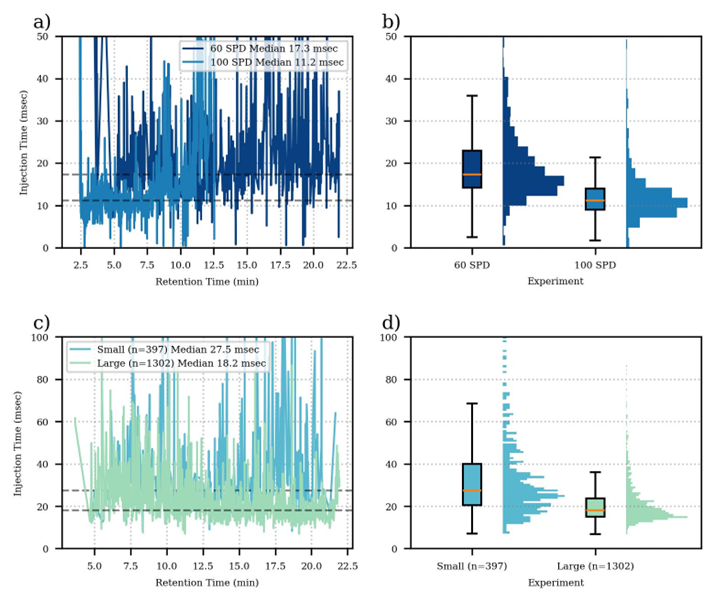

Figure S12. Injection times for the replicate experiments. a) PQ500 injection times versus retention time. b) PQ500 injection time distributions. c) E. coli injection times versus retention time d) E. coli injection time distributions.

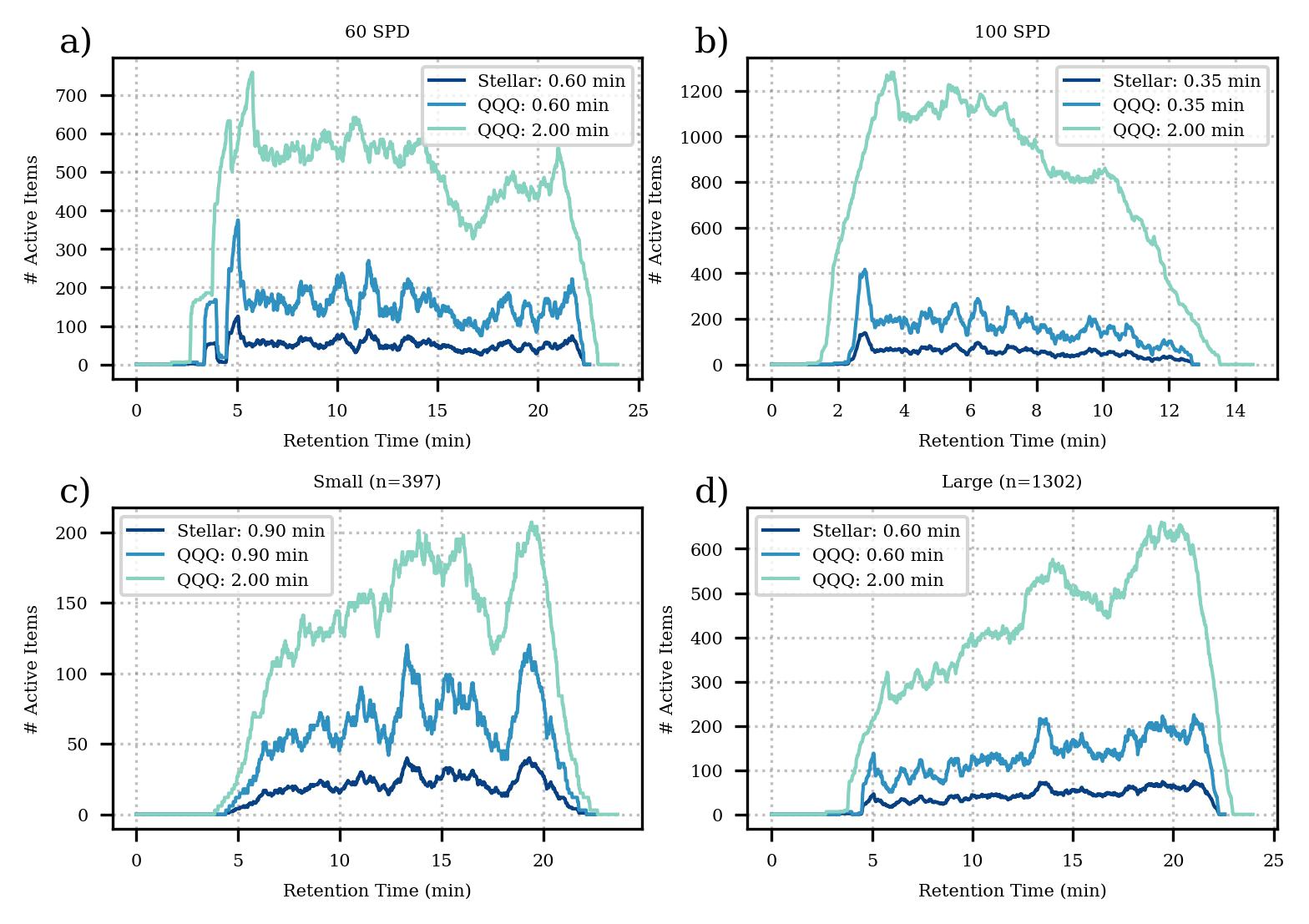

Figure S13. Concurrency of the various assays for Stellar with the scheduled acquisition times used in this study, compared to hypothetical triple quad SRM studies that used either a typical 2 minute window size, or the same size as Stellar. The triple quad concurrencies are based on 3 transitions per peptide. a) PQ500 60SPD, b) PQ500 100 SPD, c) E. coli n=397, d) E. coli n=1302.

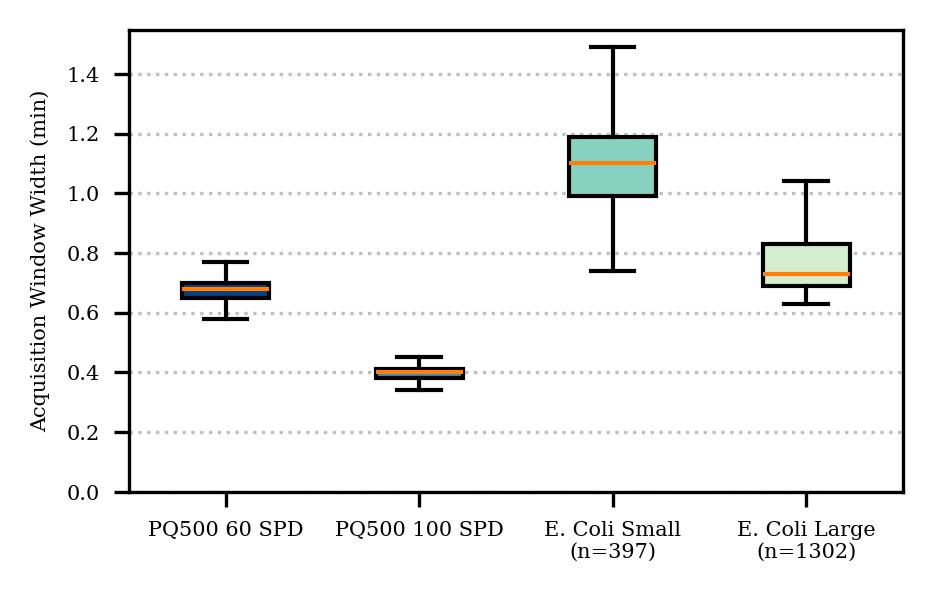

Figure S14. Distributions of actual scheduled acquisition window widths for the various experiments. Nominal widths entered into PRM Conductor were 0.6, 0.35, 0.6, and 0.6 minutes, where an optimization is applied to widen the windows where possible, and Adaptive RT adds about an LC peak width to the actually used widths.

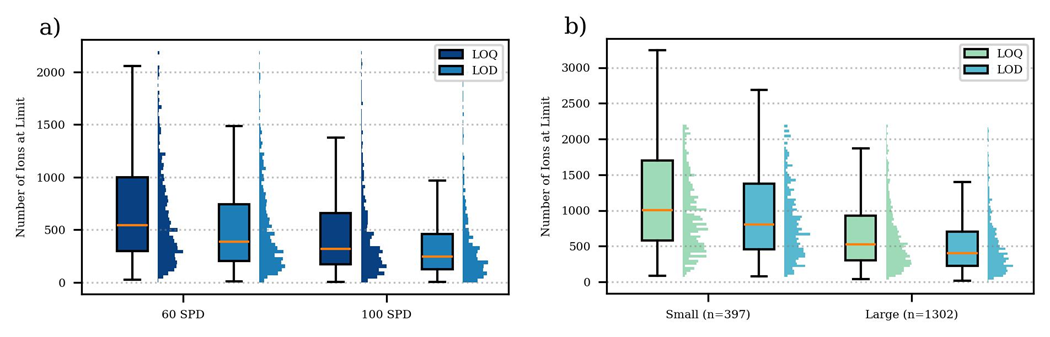

Figure S15. Distributions of number of ions used for data analysis belonging to the precursor at the limits of quantitation and detection for the various assays. a) PQ500 study, b) E. coli study.

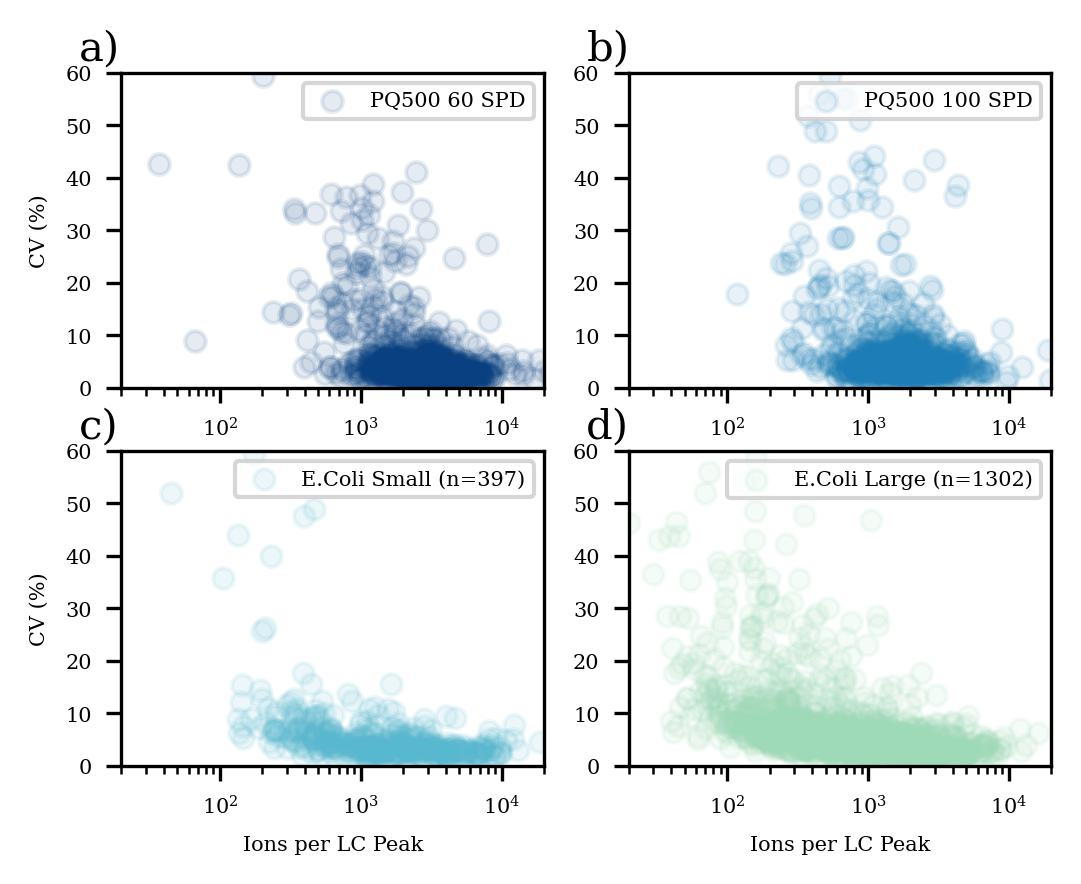

Figure S16. Coefficients of variation for LC peak areas versus the corresponding number of ions for the replicate experiments for a) PQ500 60 SPD, b) PQ500 100 SPD, c) E. coli n=397, d) E. coli n=1302.
